# Multiple-Demand Network encoding geometry balances generalization and dimensionality during novel task assembly

**DOI:** 10.64898/2026.04.22.720093

**Authors:** Ana F. Palenciano, Paula Pena, Alexandra Woolgar, Carlos González-García, María Ruz

## Abstract

On the basis of verbal instructions, humans can accomplish novel and diverse demands on the very first try. This complex phenomenon recruits structured brain activity across the frontoparietal Multiple Demand Network (MDN), which is thought to encode upcoming task parameters and guide behavior. However, it is uncertain how novel instructions are translated into efficient neural task representations. To address this, we collected functional magnetic resonance imaging (fMRI) data while participants followed a rich set of novel verbal instructions. These varied along three core dimensions: the overarching task demand (to select or to integrate stimuli information), the relevant target category (animate or inanimate items), and the visual feature that participants responded to (color or shape). We used multivariate pattern analysis (MVPA) to examine whether and how each of these dimensions was reflected in MDN activity. We contrasted two alternative representational geometries that may underpin novel task coding: low-dimensional spaces based on abstract and generalizable representations, and high-dimensional architectures hosting context-unique, conjunctive neural codes. Our results showed that anticipatory MDN activity was sensitive to the content of instructions. While selection vs. integration task demands were broadly encoded across the MDN, coding of the relevant categories and features was restricted to lateral MDN regions, namely, the intraparietal sulcus and the inferior frontal junction. Critically, the representational spaces across the MDN displayed a mixture of geometrical motifs, partially supporting our two alternative hypotheses. On the one hand, Cross-Condition Generalization Performance revealed the presence of abstract and transferable neural codes for task demand information. On the other hand, Shattering Dimensionality showed complex, high dimensional coding spaces across the MDN, structured around both task-informative and non-informative axes. Despite this, no evidence of conjunctive neural codes was observed. Overall, these findings highlight that novel instructed behavior may recruit both abstraction and high dimensionality to promote generalization while maximizing the expressivity of MDN coding spaces. More broadly, they emphasize the importance of considering encoding geometry for a computational understanding of cognitive control processes.

## Introduction

Compared with other animal species, humans show remarkable success in navigating new and versatile cognitive task environments. This adaptability is tightly linked to our employment of verbal instructions to acquire complex task procedures (Brass et al., 2017; Cole, Laurent, et al., 2013). Past research has linked this phenomenon to frontoparietal regions that assemble models or *representations* of the instructed demands (e.g., González-García et al., 2021; Muhle-Karbe et al., 2017; Palenciano et al., 2019a; Woolgar et al., 2015; Zheng et al., 2024). Building on recent research unraveling the geometry of neural representation (Badre et al., 2021), this work aims to advance our knowledge of the principles governing task coding in novel scenarios.

To effectively translate new instructions into actions, we must encode and assemble all the task parameters into a compact representation able to guide upcoming behavior (Brass et al., 2017; Cole, Laurent, et al., 2013). This process, based on cognitive control mechanisms (Norman & Shallice, 1986), has been linked to preparatory brain activity across a set of prefrontal and parietal regions (Cole et al., 2010, Hartstra et al., 2011, Palenciano et al., 2019b) that belong to the Multiple Demand Network (MDN; Assem et al., 2020, Duncan, 2010; Fedorenko et al., 2013). Using multivariate pattern analyses (MVPA), sensitive to fine-grained information coded in distributed brain activity (e.g., Haxby et al., 2014; Haynes & Rees, 2006), recent research has started to characterize the nature of novel task representations. These results have shown that different dimensions of the instructed task are encoded in advance (i.e., before stimulus presentation) within the MDN, such as targeted stimulus categories (González-García et al., 2017; Muhle-Karbe et al., 2017), relevant action plans (Palenciano et al., 2019a, 2024), or rules binding both (Cole et al., 2011; Cole, Reynolds, et al., 2013; González-García et al., 2021). The strength of these neural codes is further modulated by overarching contextual demands (González-García et al., 2021; Sobrado et al., 2022) and impacts behavioral indexes (Cole et al., 2016; González-García et al., 2021), emphasizing their role in guiding novel performance. Nonetheless, the computations that enable the fast assembly of neural task representations across the MDN remain largely unknown.

Recent models of cognitive control have stressed the role of the representational geometry for goal-oriented information coding (Badre et al., 2021; Freund et al., 2021; Musslick & Cohen, 2021). This term refers to the structure or organization that shapes how a set of features (e.g., task parameters) are encoded in a brain region’s distributed activity (Kriegeskorte & Kievit, 2013). Current proposals state that coding architectures at the service of cognitive control are fine-tuned to current demands (Badre et al., 2021; Bhandari et al., 2024; Musslick & Cohen, 2021), with two opposite geometrical motifs potentially supporting flexible task coding. On one hand, MDN regions can implement compressed representational geometries organized around a low number of task-relevant dimensions while abstracting over irrelevant information (Bernardi et al., 2020; Brincat et al., 2018). This mechanism leads to abstract and common neural codes that can easily generalize and be recombined across task contexts, a principle also known as compositional coding (Frankland & Greene, 2020; Reverberi et al., 2012; Tafazoli et al., 2026). On the other hand, some of these frontoparietal areas also display expansive, high-dimensional coding architectures, which can host segregated neural codes for each task instance (Fusi et al., 2016; Kikumoto et al., 2024; Kikumoto, Mayr, et al., 2022a, 2022b; Kikumoto, Sameshima, et al., 2022; Kikumoto & Mayr, 2020a; Rigotti et al., 2013). These geometries are thought to rely on conjunctive task coding (Hommel, 2009), which non-linearly binds all the relevant task parameters into unique activity patterns (Rigotti et al., 2013). This coding mechanism also contributes to flexible behavior by maximizing the expressivity of the encoding spaces, i.e., the number of tasks that can be differentially represented (Badre et al., 2021). Nonetheless, from a theoretical perspective, both mechanisms constitute suitable solutions for the computational challenges imposed by novel tasks, which could be coded either by abstracting and re-using a small set of common task dimensions (Cole, Laurent, et al., 2013; Cole, Reynolds, et al., 2013) or by exploiting high-dimensional coding spaces that can host a vast diversity of segregated neural task codes (Rigotti et al., 2013).

The scarce evidence to date appears to support the abstraction-based account. Instruction-driven representational geometries found in MDN regions can be decomposed into a few task dimensions (Cole, Reynolds, et al., 2013; Palenciano et al., 2019a) that generalize across contexts (Cole et al., 2011; Sobrado et al., 2022). Moreover, compositional signatures seem to be particularly relevant during the first encounters with new tasks, since experience promotes a shift toward context-specific neural patterns (Mill et al., 2025). However, the degree of abstraction might depend on the type of information being encoded. Past experiments show that generalization seems predominantly tied to higher-level task dimensions rather than lower-level features, both in novel (Pena et al., 2025) and practiced task settings (Shashidhara et al., 2024). Studies manipulating sensory modalities have further suggested that stimulus-response rule codes may not generalise between auditory and visual modes (Jackson et al. 2025). Within this complex landscape, past work has left unexplored how new task environments recruit the two geometrical features predicted by the above-mentioned models: the abstraction of the neural codes carrying task information, and the dimensionality of the underlying encoding space. Hence, a direct comparison between compositional and conjunctive novel task representation is still lacking.

To fill this gap, we employed fMRI while participants followed a rich set of novel verbal instructions. These sought to impact task processing at different hierarchical levels, manipulating the more abstract and overarching task demand (either to select a target while ignoring the other or to integrate the two targets), the allocation of attention to specific stimulus categories (either animate or inanimate targets) and the more concrete features that participants had to respond to (either the color or the shape of the frame surrounding the target). First, we used MVPA classifiers to characterize the informational content that structured activity patterns across different regions from the MDN. Second, we characterized the instruction-driven encoding spaces, focusing on two core features: abstraction and dimensionality. Lastly, motivated by recent results in the field, we examined the presence of conjunctive task representations through Representational Similarity Analysis (RSA). By doing so, we aimed to juxtapose the two main coding architectures identified in the cognitive control literature: whether novel tasks induce low-dimensional and abstract coding spaces or, alternatively, high-dimensional spaces conformed by context-specific activity patterns. These two architectures are often presented as alternatives but, as our data also show, may not necessarily be mutually exclusive.

## 1 Materials and methods

### 2.1. Participants

We collected data from 41 students from the University of Granada (Spain). All participants were native Spanish speakers, right-handed, with normal or corrected-to-normal vision, and with no history of neurological disease. In exchange for their time and effort, they received between 20€ and 25€ depending on their performance. They all signed an informed consent form approved by the Human Research Ethics Committee of the University of Granada. We excluded one participant due to technical issues with the fMRI data collection, and another one for chance-level performance (accuracy rate: 0.45), resulting in a final sample of 39 participants (25 females, 14 males; mean age = 21.6 years old, SD = 3.1 years old). The sample size was selected a priori following equivalent studies in the field (e.g. Palenciano et al., 2019a). A post hoc power analysis confirmed that 39 participants were sufficient to detect a small-medium effect size (*Cohen’s d* = 0.3) in a 2-way interaction (*see below*) with 90% statistical power (computed with the software Pangea; Westfall, 2016).

### 2.2. Apparatus, stimuli, and protocol

Each trial of our paradigm entailed a unique verbal instruction stating an “*If … then …*” conditional that referred to features of an upcoming pair of stimuli. All our instructions were in Spanish, our participants’ native language. The instructions were created by manipulating three task components that aimed to affect different information processing levels. First, the instructions indicated the trial’s overarching demand: either to select one stimulus while ignoring the other, or to integrate information from both. Second, they indicated the identities of the stimuli that should be attended to, which could be either a pair of animate (land animals and sea animals) or inanimate (tools and musical instruments) items. All instructions referred to either animate or inanimate targets, including a stimulus from each subcategory (e.g., in animate trials, a land and a sea animal were always mentioned). Third, we specified the relevant stimulus feature that participants should respond to, being either the shape or the color of the frame(s) that surrounded the target stimulus/stimuli. In selection trials, only the frame from the cued stimulus should be attended to, while in integration trials, both stimuli frames should be processed. The full set of instructions was the result of orthogonally crossing task demand (selection, integration), stimulus category (animate, inanimate), and relevant feature (color, shape), yielding eight experimental conditions. For instance, the instruction “*Atiende al martillo y a la guitarra. Si están rodeados por un rombo, pulsa L. Si no, pulsa A”,* in English: “*Pay attention to both the hammer and the guitar. If they are both surrounded by a rhombus, press L* (right finger)*. Otherwise, press A* (left finger)” belonged to the [integration, inanimate, shape] condition (see **Fig. 1** for more example instructions).

**Figure 1.**
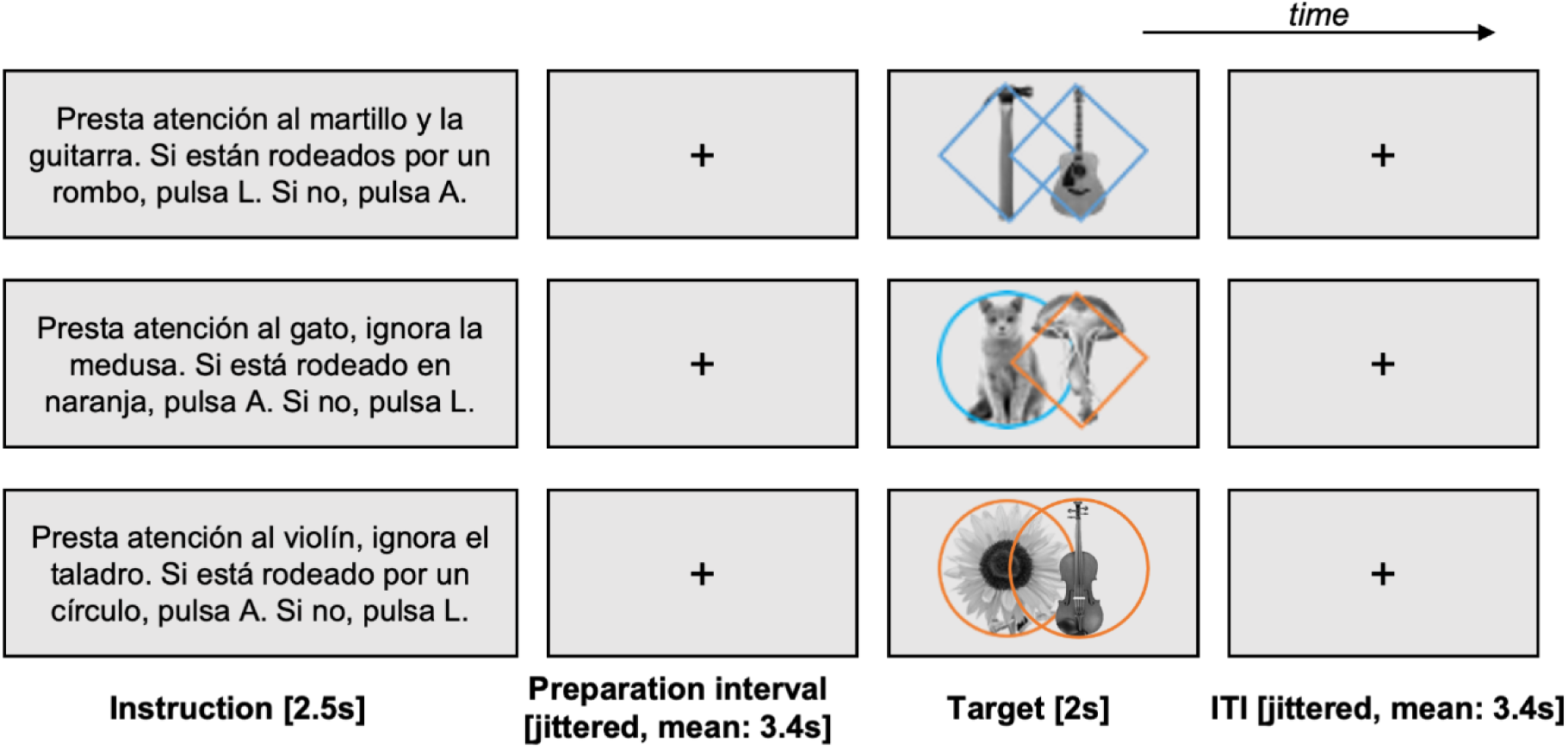
Experimental paradigm. Sequence of events across experimental conditions. Upper row. English translation: “Pay attention to both the hammer and the guitar. If they are both surrounded by a rhombus, press L. Otherwise, press A”. Regular trial from the [integration, inanimate, shape] condition. Since the stimuli satisfy the conditions of the instruction, the correct response would be a right index finger keypress (“L”). Middle row. English translation: “Pay attention to the cat, ignore the jellyfish. If it is surrounded in orange, press A. Otherwise, press L”. Regular trial from the [selection, animate, color] condition. Since the stimuli do not satisfy the conditions of the instructed statement, the correct response would be a right index finger keypress (“L”). Lower row. English translation: “Pay attention to the violin, ignore the drill. If it is surrounded by a circle, press A. Otherwise, press L”. Catch trial where a new stimulus (a sunflower) is shown instead of one of the instructed stimuli (the drill), so the correct response would be simultaneous left and right index finger keypresses (“A” and “L”). Note that in behavioral practice, participants responded with the “A” (left index) and/or “L” (right index) keys. In the scan session, however, responses were made via an fMRI-compatible button box using the left and right index fingers, corresponding to “A” and “L”, respectively. Instructions were consistent across both versions.

To increase the size and richness of our instruction pool, we additionally varied a series of task variables that were not included in our main analyses but were counterbalanced across experimental conditions. Specifically, we manipulated the identity of the target stimuli (with eight different identities: cat, dog, jellyfish, octopus, drill, hammer, guitar, and violin, all of them equally distributed across conditions), the stimulus pairings (all entailing either the two animate or the two inanimate subcategories, e.g., [cat, jellyfish]), the mapping between stimuli and task demands (e.g., in the pairing [cat, jellyfish], both stimuli were equally selected and ignored), the specific frame feature indicated in the instruction (circle or rhombus, when responding to the shape, and blue or orange, when responding to the color), and the keypresses to indicate whether the stimuli fulfilled or not the instruction (left and right index responses, linked to the keyboard keys “A” and “L”, respectively). The combination of these parameters yielded a pool of 256 instructions per participant while ensuring that the above-mentioned variables were balanced across experimental conditions. To empirically validate our experimental design, we computed the mutual information between all pairs of task parameters and compared it against a null distribution (based on 5000 permutations), ensuring statistical independence (for a similar approach, see González-García et al., 2021). Regarding the instructions’ response mapping (i.e., the response associated with the stimuli satisfying or not the instructed conditions), we kept the keypress labels “A” and “L” inside the scanner, even though an MRI-compatible joystick was employed instead of a computer keyboard. This was done to facilitate the transition between the practice (see below) and the scanner session. Please note that in Spanish, the letters “A” (left index keypress) and “L” (right index keypress) are not in conflict with the words left (“*izquierda*”) and right (“*derecha*”), as they are in English. All instructions were shown in the middle of the screen, in font Arial, size 12.

The instructions were applied to stimulus displays composed of two grayscale pictures (200 x 200 pixels) depicting animals or objects and surrounded by a colored frame. We used eight pictures (cat, dog, jellyfish, whale, screwdriver, hammer, violin, guitar) obtained from open sources with a Creative Commons license (CCBY-), and generated four versions of each, one per frame condition (rhombus-blue, rhombus-orange, circle-blue, circle-orange). The two stimuli were displayed in the midline of the screen, overlapping in 54% of their central area. Probe displays were designed so that the conditions of the instructions were satisfied in 50% of trials, equally distributing this percentage across experimental conditions. Moreover, the two frames of the probe displays could either match (e.g., if pink is the relevant feature, both probe frames are pink, or both are not pink) or mismatch (e.g., one frame is pink, the other is not pink), on the relevant dimension, independently of the task demand. The presence of congruent and incongruent probes was also balanced across experimental conditions. Finally, we pseudo-randomized the location of the two probes on the screen (left or right side).

The sequence of events within each trial is shown in **Fig. 1**. First, an instruction was shown in the center of the screen for 2.5 s, followed by a preparation interval with a jittered duration (mean: 3.4s, range: [2.6-5.8] extracted from a pseudo-logarithmic distribution, e.g., Palenciano et al., 2024). Then, the probe stimuli appeared on the screen for 2s. In 87.5% of (regular) trials, the two stimuli corresponded to the identities stated in the instruction, and participants responded according to their frame color or shape. However, in 12.5% of trials (catch, an example is shown in the lower row from **Fig. 1**), one of the two stimuli was an object that had not been mentioned in the instruction (either a vehicle or a plant). In these cases, participants had to respond with both index fingers simultaneously. Catch trials sought to ensure that both probe identities were processed during the instruction encoding (and not merely the attended stimulus in selection trials, or none in integration trials). After a second jittered interval (same properties as above), the next trial began. Participants completed a total of 256 trials, divided into 8 runs of 32 trials each.

Before the MRI, participants completed a practice session to familiarize themselves with the paradigm, using an independent set of instructions that were not employed in the experimental blocks. A minimum of 80% of correct responses was required to continue to the scanning session. In the scanner, participants completed eight runs of the main paradigm. Afterwards, during a final functional run, participants completed a localizer 1-back task. This showed the same eight stimuli individually, without the frame and in a pseudo-random order, and participants responded if the same sub-category (i.e., land animals, sea animals, tools, music instruments) was repeated in two consecutive trials. Data from this localizer task were not analyzed in the current work, and hence, it is not discussed further.

### 2.3. Experimental design and behavioral analyses

Our paradigm followed a within-subject design, with three factors of interest: (1) task demand (to select or to integrate probe information), (2) target category (animate and inanimate probes), and (3) relevant feature to respond to (shape or color of the frame). These three variables were fully crossed, resulting in eight experimental conditions. Overall, we collected 32 observations per cell of this design (four per fMRI run).

For the behavioral analyses, we discarded data from catch trials and those with RT above and below 2 standard deviations from the participant’s mean. For the reaction times (RT) analysis, we further excluded error trials. Then, we addressed the effect of our three factors on accuracy rate and RT in two independent repeated measures ANOVAs (rmANOVAs).

### 2.4. MRI data acquisition and preprocessing

fMRI data was acquired with a 3T Siemens Magnetom Prisma Fit scanner located at the Mind, Brain, and Behavior Research Center (University of Granada, Spain), using a 64-channel head coil. The following sequences were acquired, in order: a T2-weighted anatomical image, a phase and magnitude field map images, and nine T2* EPI functional runs (TR = 1730 ms, TE = 30 ms, 50 slices per volume, flip angle = 66°, FOV = 210 mm, voxel size = 2.5 mm^3^, distance factor = 0%, acceleration factor = 2, slice orientation following the participants’ AC-PC line). The first eight runs comprised 212 volumes each and were collected during the main experimental paradigm. The last run, with 252 volumes, covered an additional localizer task. Finally, we obtained a high-resolution anatomical T1-weighted image using an MP-RAGE sequence (TR = 2250 ms, TE = 4.18 ms, 176 slices, flip angle = 9°, FOV = 256 mm, voxel size = 1 mm^3^). In total, the scanning session lasted ∼72 minutes.

Data were preprocessed with SPM12 (Penny et al., 2006). The functional volumes were corrected for magnetic field inhomogeneities (using the field mapping images), spatially realigned and unwarped to account for motion artifacts (using the runs’ first volume as reference) and slice-time corrected (using the middle slice as reference). Then, we deidentified (i.e., defaced) the T1 anatomical images, co-registered them to the functional volumes, and carried out a segmentation. To avoid distorting the fine-grained multivoxel activity patterns, we performed our main multivariate analyses using the (non-normalized and non-smoothed) functional volumes. However, for the univariate control analyses, the EPI images were further normalized to the MNI space using the transformation matrices provided by the T1 segmentation and then smoothed with an 8mm FWHM Gaussian kernel. All MRI data were organized according to the BIDS standard (Gorgolewski et al., 2016).

### 2.5. fMRI data analyses

We estimated a series of first-level GLMs on each participant’s data. These included regressors covering the trials’ instruction, preparation and target stages, separately for the eight experimental conditions (i.e., combinations of Task Demand, Target Category and Relevant feature). All regressors were modeled using a stick function (i.e., 0s duration) convoluted with the SPM’s canonical hemodynamic response function (HRF). An assessment of the collinearity among the three task events regressors using Pearson’s correlation confirmed that the GLM’s coefficients were estimable (*r_Instruction-Preparation_* = 0.70, *r_Instruction-Target_* = 0.13, *r_Preparation-Target_* = 0.52). Additional nuisance regressors included error trials (locked to the instruction onset and covering the full trial duration) and the six movement parameters computed during the realignment. The estimation followed a traditional LSU (Least-Squares Unitary) approach (i.e., beta per run). These GLMs were first estimated with smoothed and normalized volumes to identify areas univariately engaged by our experimental paradigm. This was done with one-sample *t*-tests that contrasted each trial event against the implicit baseline. Then, we estimated the GLMs using unnormalized and unsmoothed images to perform the multivariate pattern analyses. This provided a beta weight brain map per event, condition, and MRI run. Finally, we also contrasted these images against the implicit baseline (using one-sample *t*-tests), obtaining brain-wise *T* maps where the noise was normalized in a univariate fashion (Misaki et al., 2010). These were used for the main multivariate analyses.

In addition, we estimated a series of trial-wise GLM models to obtain separate *T* maps for each instruction employed in our task. This allowed us to explore finer-grained information provided by the instruction pool (see section 2.5.5). To reduce collinearity, we employed an LSS (Least-Squares Separate) approach (Arco et al., 2018; Mumford et al., 2012, 2014), computing an individual GLM per trial with the following regressors: (1) the preparation interval of the targeted trial, (2) the preparation interval from the remaining trials, (3) the instruction stage from all trials, and (4) the probe stage from all trials. Additional nuisance regressors also modeled error trials and the six movement parameters. Only unnormalized and unsmoothed volumes were employed to estimate the models. Again, we contrasted the resulting beta weight maps against the implicit baseline to obtain trial-wise T maps.

MVPA was implemented with The Decoding Toolbox (Hebart et al., 2014) and custom-made MATLAB scripts. We generated the ROI results plots with the Python library Seaborn (Waskom, 2021) and whole-brain maps using MRIcroGL (https://www.nitrc.org/projects/mricrogl).

#### 2.5.1. ROI selection

Our planned analyses were performed on a series of MDN ROIs, defined based on the work of Fedorenko and colleagues (2013) and using the templates available online (http://imaging.mrc-cbu.cam.ac.uk/imaging/MDsystem). We employed nine ROIs (**Fig. 2A**) that covered the pre-supplementary motor area (preSMA) and anterior cingulate cortex (ACC; referred to as preSMA/ACC ROI), and left and right inferior frontal sulcus (L_IFS, R_IFS), inferior frontal junction (L_IFJ, R_IFJ), anterior insula-frontal operculum area (L_aI/fO, R_aI/fO) and the intraparietal sulcus (L_IPS, R_IPS)t. The ROI masks, defined in the standard MNI space, were co-registered and inverse-normalized to each participant’s anatomical space using the T1 volume and later binarized.

**Figure 2.**
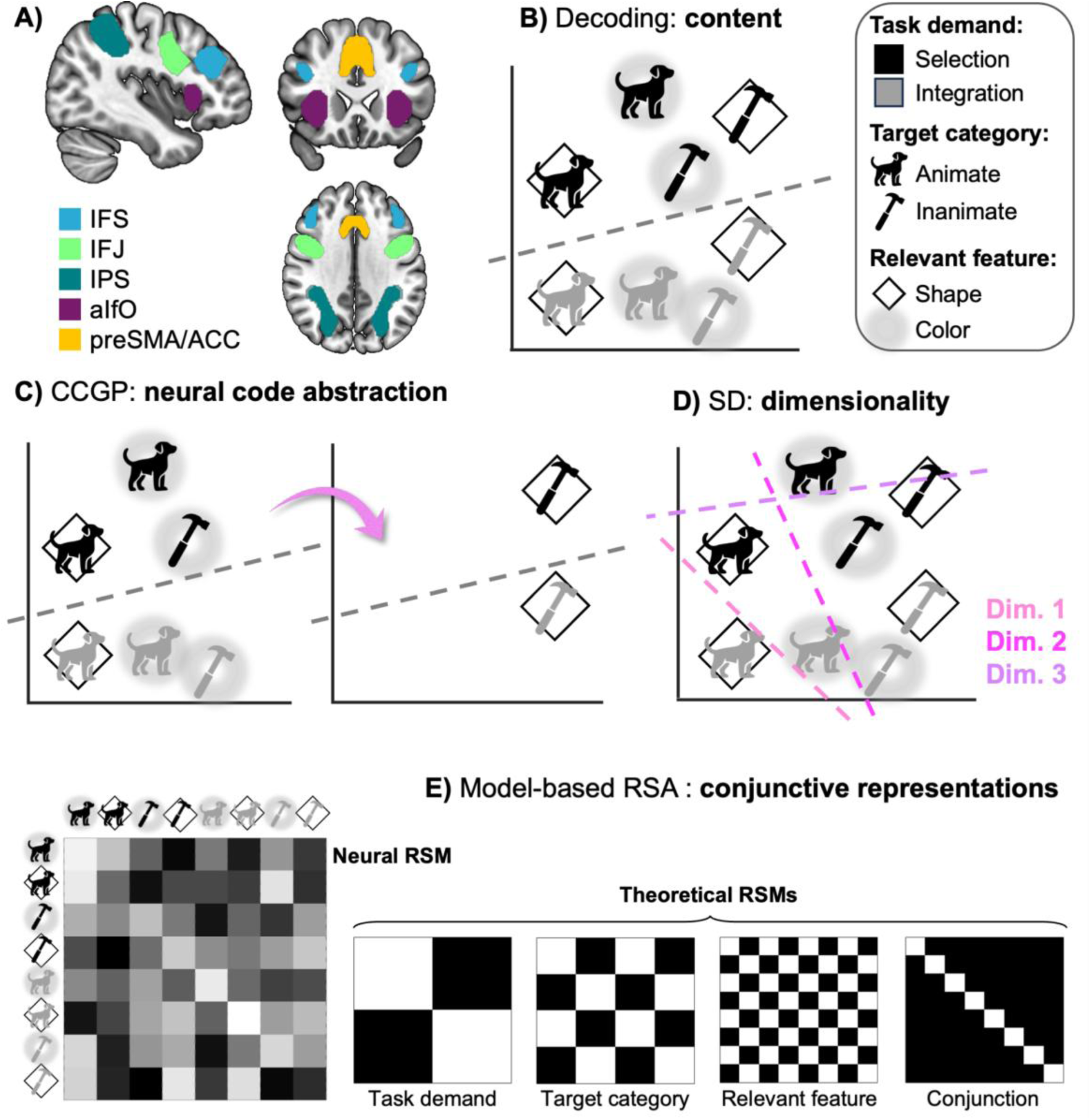
Multivariate analysis pipeline. **A)** MDN ROIs. **B)** Decoding classifiers were used to identify information content. The figure illustrates the conditions’ activity patterns against a simplified coding space, with the grey line depicting a linear classifier decoding task demand (i.e., differentiating selection from integration trials). The inset shows the symbols used in **B-E** to represent the eight experimental conditions/contexts. **C)** CCGP was used to quantify neural code abstraction. The example shows one iteration of the cross-decoding procedure, where a classifier is trained to decode selection from integration trials in a subset of contexts and later tested (pink arrow) in the left-out contexts. **D)** SD was used to estimate the dimensionality of the ROIs’ encoding spaces. Above-chance classifications decoding subsets of interpretable or non-interpretable task contexts (hyperplanes shown in different pink hues) were interpreted as independent coding dimensions. **E)** Model-based RSA was used to identify conjunctive task representations. The figure illustrates (left) a neural RSM reflecting the similarity among activity patterns from all the pairwise context combinations (white: maximum similarity, black: minimum similarity). (Right) Multiple regressions were fitted to predict the ROIs’ RSM with models based on the three task components and their conjunction. The conjunctive model predicts minimal distance for same-context patterns and maximal distance for the remaining combination.

#### 2.5.2. Decoding analysis

First, machine learning classifiers were used to detect information coded in MDN during novel instruction preparation. Focusing on the activity patterns from the preparation interval, we applied standard decoding analyses to classify them according to the three task components manipulated in the instructions: the relevant task demand, the targeted stimulus category, and the feature to respond to. To do so, linear Support Vector Machine (SVM) algorithms were trained and tested to classify the two levels of each variable (i.e., selection vs. integration, animate vs. inanimate, shape vs. color, respectively) using data from each ROI (**Fig. 2B**). The ROIs’ activity patterns consisted of *T*-value vectors obtained from the above-mentioned LSU (i.e., beta per run) GLM. A leave-one-run-out cross-validated scheme was applied, where the classifier was trained with data from seven out of the eight fMRI runs and later tested against the remaining run. We iterated this procedure until all runs were used as a testing dataset once. Balanced accuracy (minus the chance level, i.e., 0.5) was computed for each cross-validation fold and later averaged. The employment of the same number of activity vectors (i.e., exemplars) per condition within each classification ensured that findings were not biased towards a preferred class. Then, at the group level, we addressed whether the subjects’ mean balanced accuracy was greater than zero using Wilcoxon one-sided signed-rank tests. Within each task component, a Bonferroni-Holm multiple comparison correction (Holm, 1979) was applied to control for the number of ROIs analyzed. We complemented those findings with Bayesian one-sample T-tests to provide evidence regarding the null hypothesis, in this context, chance-level decoding performance. Interpretation of Bayes Factor was done following standard practices; i.e., 0.3 < BF_01_ < 3 as inconclusive evidence for the null effect, BF_01_ > 3 as moderate evidence for the null effect, BF_01_ > 10 as strong evidence for the null effect (Schmalz et al., 2023). All statistical inference in the multivariate analysis was performed using the same approach.

To explore data outside the MDN and ensure the transparency of our results, we also conducted an exploratory whole-brain analysis. For each task component, we applied the same decoding analysis using a searchlight procedure. The classification was performed using activity patterns from a 5-voxel radius sphere that iterated through every location of the brain, assigning the decoding output to the sphere’s central voxel. Group statistics were assessed with a one-sample t-test comparing the balanced accuracy minus chance against zero. The results were corrected for multiple comparisons using an FWE criterion at *p* < .05 (from an initial threshold of *p* < .001 and k = 10).

#### 2.5.3. Cross-Condition Generalization Performance

Next, we aimed to address whether the three task components were encoded in an abstract, generalizable format. Cross-Condition Generalization Performance (CCGP) was used to summarize how well a given variable’s activity patterns generalize or transfer between conditions, which has been interpreted in the past as a signature of neural code abstraction (Bernardi et al., 2020).

The CCGP index is estimated by applying an exhaustive protocol of cross-decoding classifiers that are trained and tested in separate task contexts (Bernardi et al., 2020; Bhandari et al., 2024, **Fig. 2C**). In our study, these contexts corresponded to the eight experimental conditions resulting from crossing the three main manipulations. Depending on the task component analyzed, these were arranged into two groups, one per level of manipulation. For instance, to compute the task demand CCGP, one set of conditions included the four selection contexts (such as [selection, animate, color]), and the other one, the remaining four integration contexts (such as [integration, inanimate, shape]). Each CCGP was estimated by training SVM classifiers with data from six out of the eight contexts, ensuring a balanced presence of the two levels of the targeted task component (e.g., three selection and three integration contexts). Then, the classifier performance was tested using the remaining two contexts, one per component level (e.g., the held-out selection and integration context). With our experimental design, for each task component we could sort these contexts into 16 different cross-classification schemes. Specifically, the four contexts of each component level could be arranged into (three) training and (one) testing dataset in four different ways ((4 3) = 4). Crossing the four arrangements of the two components’ levels led to the 16 interactions. For each component and ROI, we trained and tested the 16 cross-decoding SVM classifiers and averaged the resulting balanced accuracy rates (minus chance), which corresponded to the CCGP score. Group-level inference was performed on this index and in the output of each of the cross-decoding iterations to ensure that significantly above-chance CCGP scores were not driven by particularly high accuracies in reduced sets of condition arrangements.

Additionally, we implemented exploratory whole-brain analyses, computing the CCGP for each task component with a searchlight procedure. For that, we used the protocol described in the *Decoding analysis* section, but now applying the CCGP classification scheme within each searchlight sphere.

Finally, it is worth noting that the CCGP was computed by cross-decoding classifiers across contexts, but not cross-validating them against independent MRI runs’ data, as we did in the standard decoding analyses. Although unlikely, there is a chance that these measurements were biased by run-specific noise captured by the classifiers. To rule out this possibility, we complemented our CCGP results with a more restrictive control analysis that inserted a leave-one-run-out cross-validation procedure within each cross-decoding scheme. This way, the cross-decoding classifiers were trained and tested with data from independent task contexts and MRI runs.

#### 2.5.4. Shattering Dimensionality

We also aimed to estimate the dimensionality of the overarching representational space of each MDN ROI. To do so, we computed the Shattering Dimensionality (SD), a measurement based on binary classifiers that approximates the minimum number of axes structuring a region’s distributed activity (Bernardi et al., 2020; Rigotti et al., 2013). This minimum dimensionality is identified, from this approach, with the maximum number of significant, above-chance binary classifications that can be performed in a given dataset (**Fig. 2D**). Hence, this index informs on whether the MDN ROIs display low-dimensional coding spaces constrained by the three task dimensions manipulated or show, instead, more expansive high-dimensional structures.

To compute the SD, we first grouped the eight experimental conditions into all possible balanced dichotomies of four vs. four conditions, regardless of their interpretability (Bernardi et al., 2020; Rigotti et al., 2013). Our design provided an overall of 35 dichotomies (i.e., (8 4) = 70, which divided by 2 to omit duplicates led to the 35 dichotomies), with three of them corresponding to the manipulated task components and the remaining 32 being non-informative regarding our paradigm. With this method, 35 was the dimensionality ceiling that could be estimated. Separately for each ROI, we trained and tested linear cross-validated classifiers to decode each one of these dichotomies. In our results, we reported the total number of significant above-chance classifications per ROI as the SD-estimated dimensionality.

Last, to explore in greater detail the nature of the coding dimensions identified through the SD protocol, we additionally computed the CCGP score of the full set of 35 dichotomies. To do so, we extended the procedure described in the previous section to the non-informative binary dichotomies identified in the SD analysis (see Bernardi et al., 2020 for a similar approach).

#### 2.5.5. Model-based Representational Similarity Analysis

The analyses described so far treat each task component independently, leaving unaddressed the relationship among them during their assembly into coherent control representations. In this regard, recent studies have emphasized that brain responses to different task parameters are non-linearly combined into conjunctive neural codes (Kikumoto et al., 2024; Kikumoto, Mayr, et al., 2022; Kikumoto, Sameshima, et al., 2022; Kikumoto & Mayr, 2020). Following this research, we employed model-based Representational Similarity Analysis (RSA) to address whether the three instructed task components were jointly coded in such a conjunctive fashion.

RSA (**Fig. 2E**) estimates the ROIs’ encoding space by computing the similarity between activity patterns from different experimental conditions, conveying these measurements into representational similarity matrices (RSM). We employed the cross-validated Pearson coefficient as a similarity metric, a robust index that is insensitive to differences in signal mean or variance between conditions, avoids positive biases, and provides interpretable values (Ritchie et al., 2021; Walther et al., 2016). This measurement was computed with a leave-one-run-out cross-validation procedure. Specifically, in each iteration we averaged the condition-wise activity patterns from seven out of the eight fMRI runs, providing a single mean pattern per condition, which was Pearson-correlated with the left-out run’s activity patterns. That provided eight-by-eight RSMs capturing the correlation between all the pairwise combinations of our eight experimental conditions. The cross-validation led to eight RSMs, which were then averaged into a single matrix per ROI. These cross-validated RSMs were non-symmetrical, and their diagonal contained interpretable values, allowing us to include the full matrices in the analyses.

Next, we decomposed the neural RSMs using a series of theoretical models, which expressed predicted between-condition similarity. Specifically, we used four categorical models: (1) task demand, (2) target category, (3) relevant feature, and (4) their conjunction. The first three were matrices where pairs of conditions sharing the same level of the corresponding component were labeled with ones (i.e., maximum similarity) and those featuring different levels, with zeros (i.e., minimum similarity). For instance, in the task demand RSM, the conditions [selection, animate, shape] and [selection, inanimate, color] were expected to be maximally similar and coded with a one, while the pair [selection, animate, shape] and [integration, animate, shape] was expected to be maximally dissimilar and coded with a zero. Replicating the strategy followed in previous studies (e.g., Kikumoto & Mayr, 2020), the conjunction model was generated after multiplying the first three, which resulted in an RSM that contained a diagonal of ones and zeros elsewhere, predicting condition-unique activity patterns. Pearson’s correlation assessment of collinearity among the theoretical RDM models revealed full orthogonality between the task demand, target category, and relevant feature RDMs, and a moderate correlation of r = 0.38 between each of these models and the conjunctive RDM. Last, we fitted linear multiple regressions predicting the ROIs’ empirical RSMs with the four theoretical RSMs as predictors. The regressions provided four beta coefficients per ROI and participant, reflecting how well each model captured the coding structure.

Finally, and in an exploratory fashion, we ran an additional model-based RSA focusing on the trial-wise activity patterns obtained from the LSS GLMs estimation. This was done to account for potential conjunctive representations that mixed lower-level instruction parameters beyond the primary task components. Our main analyses reported so far targeted the three broader task components that our experimental design was tailored for. However, our pool of instructions varied along a diverse set of parameters that may have also constrained the task coding geometry, such as the specific frame feature that participants had to detect, the response set required in each trial, or their conjunction. After estimating trial-by-trial LSS GLM models, we repeated the model-based RSA with the following modifications. First, we built trial-wise empirical RSMs, conveying the relationship between pairs of individual instructions. Second, to retain this variability, we avoided the cross-validation procedure, computing the z-score Pearson correlation coefficients between each pair of activity patterns. We retained the lower triangle from these symmetrical RSMs, excluding diagonal values. And third, in addition to the task demand, target category, and relevant feature described above, we included three more theoretical RSMs to the multiple regression, coding the specific shape or color that participants responded to (orange, blue, circle, rhombus frames), the instructed response mapping (associating a right or a left response to the conditions of the instruction being met by the stimuli), and the conjunction between these lower-level task parameters. The remaining analysis parameters remained identical to the main RSA.

## 3. Results

### 3.1. Behavioral results

Before the behavioral analyses, we filtered out abnormally fast or slow trials (RTs of ± 2SD from the participant’s mean), excluding an average of 4.3% trials per participant (range: 0.8-9.4%). On average, participants responded correctly in 90.5% of regular trials (SD = 5.7%), with a mean RT of 1085ms (SD = 171ms), and committed 2.5% (SD = 3.0%) of omission errors. Hence, despite the demanding nature of our paradigm, performance was high and in line with studies employing similar paradigms (e.g., Palenciano et al., 2019a; Pena et al., 2025). In catch trials, participants responded with a mean accuracy rate of 94.9% (SD = 4.6%) and RT of 907ms (SD = 93ms). Catch trial performance was equally accurate under selection and integration demands (T_38_ = 1.13, *p* = .268, selection: M = 94.1%, integration: M = 95.7%), although faster in the latter (T_38_ = 2,85, *p* = .007, selection: M = 935ms, integration: M = 903ms). These data indicate that the category of the two targets was processed under both demand conditions.

We analyzed the impact of the three task components on accuracy and reaction times through two separate rmANOVAs. In accuracy rates (**Fig. 3, left**), we found a significant main effect of Task Demand (F_1,38_ = 18,44, *p* < .001, η_p_^2^ = 0.33), with more accurate responses to integration (M = 92.3%, SD = 5.0%) than selection demands (M = 88.6%, SD = 7.4%), and of Target Category (F_1,38_ = 18,16, *p* < .001, η_p_^2^ = 0.32), with higher accuracy to animate (M = 91.0%, SD = 5.3%) than inanimate targets (M = 89.9%, SD = 6.7%). No significant interactions were found (all *p*s > .1). In RT data (**Fig. 3, right**), we found significant main effects of Task Demand (F_1,38_ = 52.28, *p* < .001, η_p_^2^ = 0.58), with faster responses in integration (M = 1032ms, SD = 174ms) than selection trials (M = 1141ms, SD = 182ms), of Target Category, (F_1,38_ = 97.01, *p* < .001, η_p_^2^ = 0.72), with shorter RTs to animate (M = 1069ms, SD = 165ms) than inanimate targets (M = 1100ms, SD = 179ms), and of Relevant Feature, with faster performance on color (M = 1056ms, SD = 172ms) than shape (M = 1114ms, SD = 172ms) judgments. We also found a significant interaction between Task Demand and Relevant Feature (F_1,38_ = 12.85, *p* < .001, η_p_^2^ = 0.25), driven by differences between color and shape trials under selection demands (T_38_ = 6.05, *p* < .001, Cohen’s d = .28; color: M = 1117ms, shape: M = 1167ms) but not in the integration condition (T_38_ = 1.66, *p* = .102, Cohen’s d = .08; color: M = 1026ms, shape: M = 1040ms).

**Figure 3.**
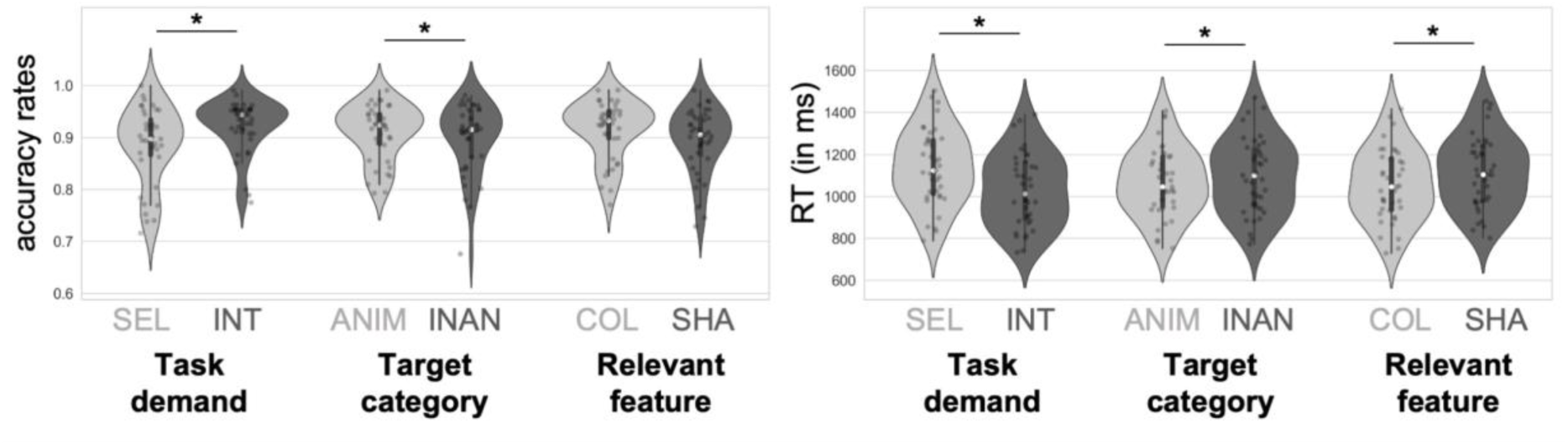
Behavioral results. Violin plots depicting the subject-wise averaged accuracy rates (left) and RTs (right) across task components and experimental conditions. Gray dots indicate individual subject means, and the white dot shows the sample median. Thick vertical lines indicate the interquartile range (Q1-Q3). Asterisks indicate significant differences between conditions’ means (i.e., significant main effects from the repeated measures ANOVA). *SEL: selection, INT: integration, ANIM: animate, INAN: inanimate, COL: color, SHA: shape*.

### 3.2. fMRI results

As a sanity check, we first ran a series of univariate one-sample *T*-tests contrasting the beta maps linked to the encoding, preparation, and execution of instructions against the implicit baseline. These results, detailed in **Supp. Fig. 1** and **Supp. Table 1** showed that our instruction-following paradigm generated univariate signal increases along the MDN, as well as in specialized perceptual and action-oriented systems, the basal ganglia, and the thalamus. While the significant clusters were substantially more widespread during instruction encoding and execution, we also detected heightened MDN activation during the preparation interval, in line with past results (González-García et al., 2017; Muhle-Karbe et al., 2017; Palenciano et al., 2019a).

Next, we applied a multivariate pipeline to characterize the content and geometry shaping MDN activity. All these analyses focused on activity patterns recruited during the preparation interval.

#### 3.2.1. Informational content

As a first step, we aimed to identify which MDN regions encoded the different task components manipulated across the instructions. Our findings (**Fig. 4A** and **Table 1)** evidenced that the prospective task demand (i.e., to select or to integrate stimulus information) was broadly encoded across the MDN. The classifiers’ balanced accuracy rates were significantly above chance in all the ROIs explored, including the preSMA/ACC and the bilateral IFS, IFJ, and IPS. The classifiers’ mean accuracies numerically peaked in the left and right IPS and the preSMA/ACC, followed by the bilateral IFJ. Conversely, the category of the upcoming targets was only decodable from the right IPS activity patterns. In the remaining regions, the classifier performance did not differ from chance level. Bayesian one-sample T-tests (see **Table 1**) provided moderate evidence supporting the null effect (i.e., chance-level accuracy) in most of these regions, except the L_IPS and the R_IFJ, where this test went in the same direction although providing more uncertain results. Finally, the relevant feature of the stimuli (either the color or the shape of the frame surrounding the targets) could be significantly decoded in the L_IFJ and the bilateral IPS. A similar trend was found in the left IFS, although this test did not survive the multiple comparison correction. Again, after conducting further Bayesian T-tests, we found moderate evidence supporting chance-level accuracy in the remaining regions, except in the R_IFJ, where evidence was inconclusive.

**Figure 4.**
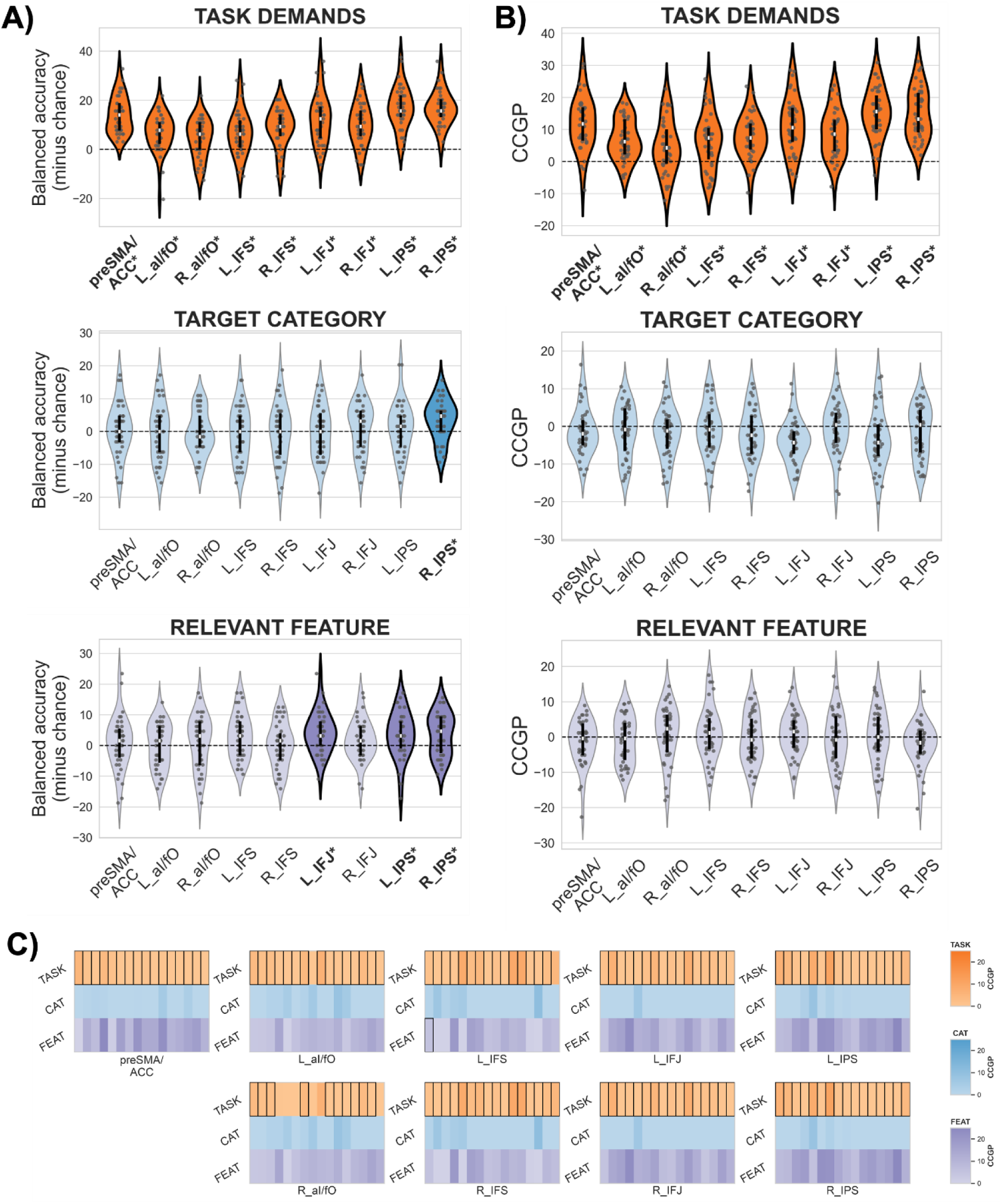
Neural coding of the three task components: decoding and Cross-Condition generalization performance (CCGP) results. **(A)** Violin plots depicting cross-validated balanced accuracy minus chance when decoding the task demand (upper figure), the target category (middle figure), and the relevant feature (lower figure) in each of the ROIs. Grey dots show individual participant means. The sample median is shown within the violin as a white dot. Significant above-chance decoding is indicated with thicker violin plot outlines and darker violin plot colors, and asterisks and bold font in the ROI labels. **(B)** Violin plots depicting the ROIs CCGP scores, using the same plotting conventions as in (A)**. (C)** CCGP across cross-decoding interactions for each ROI. Each ROI matrix depicts the three task components in rows (from top to bottom: task demand, target category, and relevant feature) and the 16 cross-decoding iterations in columns. Black frames around cells indicate significantly above-chance cross-classifications (corrected for multiple comparisons). The color intensity shows the specific CCGP value of that cell, anchored uniformly across all regions and colormaps. CCGP values at or below chance (≤ 0) are depicted with the baseline light intensity of the respective colormap

**Table 1.**
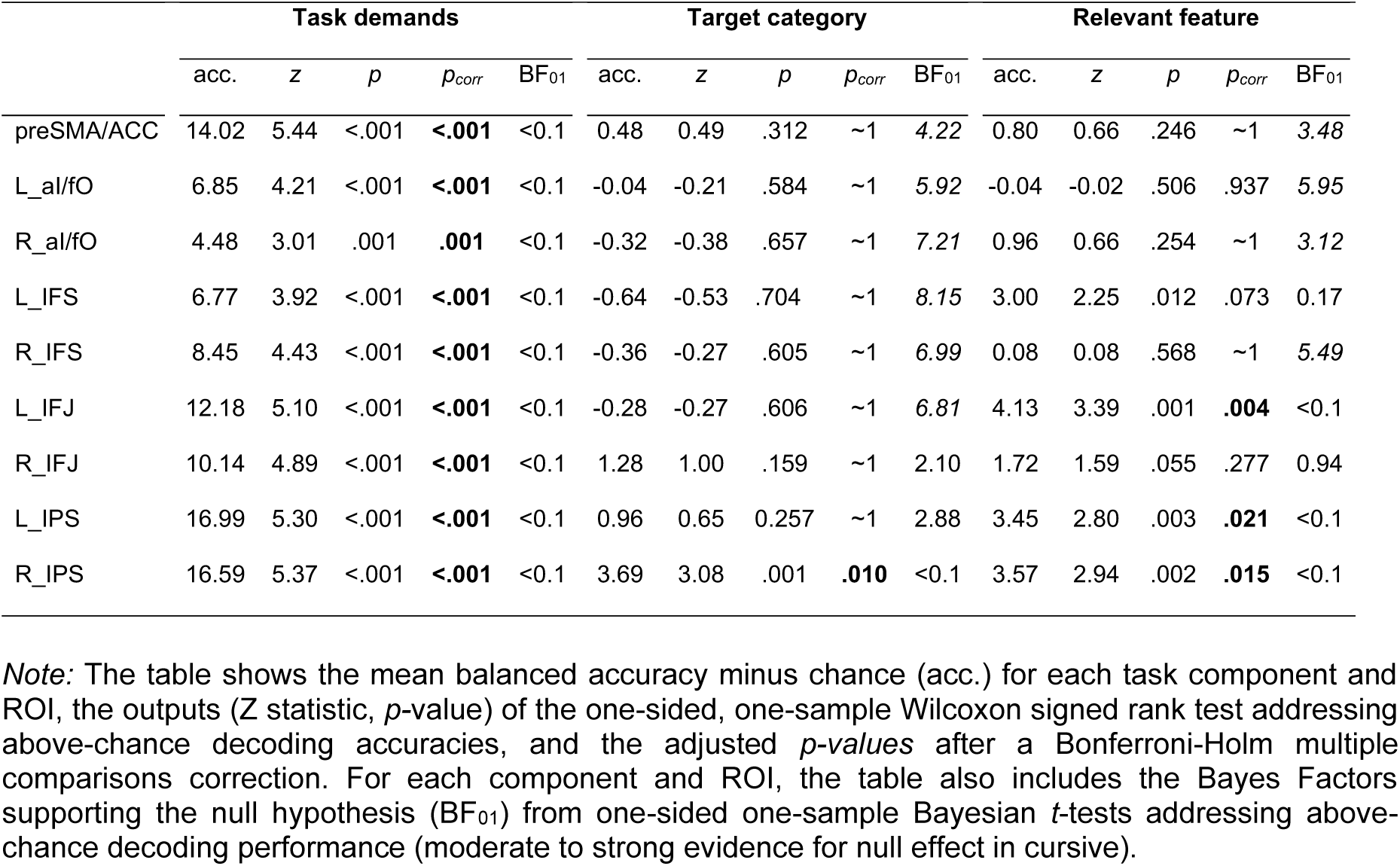
Results from cross-validated classifiers decoding the three task components across ROIs.

To complement the ROI analyses, we conducted exploratory whole-brain decoding analyses for the three task components (**Supp. Fig. 2** and **Supp. Table 2**). This analysis confirmed that task demand could be decoded within the MDN, but also broadly outside this system, including the medial prefrontal cortex, the precuneus, the temporal cortex, and specialized visual and motor brain regions. The results from the relevant feature component showed significant decoding confined to the left middle and inferior temporal gyri, and the primary and secondary visual cortex. No clusters survived FWE multiple comparison correction for Target Category.

#### 3.2.2. Neural code abstraction

Next, we focused on the format underlying these task-related representations and estimated the CCGP, an index of neural code abstraction that measured how well the activity patterns generalized across contexts. By applying an exhaustive cross-decoding protocol, we computed a CCGP score for each ROI and task component.

These results (**Fig. 4B** and **Table 2**) showed that the task demand component was systematically encoded in an abstract and generalizable fashion. This effect was found in all the MDN ROIs. Numerically, the neural code abstraction was highest in the bilateral IPS, followed by the preSMA/ACC and the L_IFJ. To rule out the possibility that these findings were driven by a subset of specific cross-decoding iterations, we also performed a more detailed examination across the different arrangements of task contexts employed to train and test the classifiers (see **Fig. 4C**). In most of the regions, the classifiers significantly cross-decoded the selection/integration demands across all the contexts arrangements (preSMA/ACC, R_IFS, L_IFJ, R_IFJ, L_IPS, R_IPS), or in 15 out of 16 arrangements (L_aI/fO, L_IFS), with the only exception of the R_aI/fO, where the test was significant in 10 out of 16 arrangements. These findings affirm that the mean CCGP scores provided a reliable insight into the cross-decoding algorithm performance.

**Table 2.** Results from Cross-Context Generalization Performance (CCGP) analysis, estimating the abstraction of activity patterns encoding the three task components across ROIs.

|  | Task demands |  |  |  |  | Target category |  |  |  |  | Relevant feature |  |  |  |  |
| --- | --- | --- | --- | --- | --- | --- | --- | --- | --- | --- | --- | --- | --- | --- | --- |
|  | CCGP | *z* | *p* | *p<sub>corr</sub>* | BF<sub>01</sub> | CCGP | *z* | *p* | *p<sub>corr</sub>* | BF<sub>01</sub> | CCGP | *z* | *p* | *p<sub>corr</sub>* | BF<sub>01</sub> |
| preSMA/ACC | 11.92 | 5.09 | <.001 | **<.001** | <0.1 | -1.23 | -1.63 | .948 | ~1<sup>a</sup> | 11.93 | -1.44 | -0.86 | .812 | ~1 | 11.72 |
| L_al/fO | 6.80 | 4.74 | <.001 | **<.001** | <0.1 | -1.29 | -0.96 | .831 | ~1<sup>a</sup> | 11.66 | -0.90 | -0.89 | .812 | ~1 | 11.34 |
| R_al/fO | 4.51 | 2.95 | .002 | **.002** | <0.1 | -1.85 | -1.61 | .947 | ~1<sup>a</sup> | 11.84 | 0.94 | 1.05 | .146 | .806 | 2.31 |
| L_IFS | 6.42 | 3.88 | <.001 | **<.001** | <0.1 | -1.04 | -0.89 | .814 | ~1<sup>a</sup> | 9.85 | 1.15 | 0.76 | .223 | ~1 | 2.38 |
| R_IFS | 7.67 | 4.81 | <.001 | **<.001** | <0.1 | -2.08 | -1.72 | .957 | ~1<sup>a</sup> | 13.91 | -0.56 | 0.23 | .409 | ~1 | 4.92 |
| L_IFJ | 11.25 | 5.13 | <.001 | **<.001** | <0.1 | -3.94 | -3.75 | .999 | ~1<sup>a</sup> | 22.92 | 0.86 | 0.91 | .180 | ~1 | 3.08 |
| R_IFJ | 8.75 | 5.00 | <.001 | **<.001** | <0.1 | -0.53 | -0.31 | .621 | ~1<sup>a</sup> | 6.75 | -0.57 | -0.51 | .693 | ~1 | 10.06 |
| L_IPS | 14.99 | 5.39 | <.001 | **<.001** | <0.1 | -3.47 | -2.57 | .995 | ~1<sup>a</sup> | 18.78 | 0.04 | 0.11 | .458 | ~1 | 6.08 |
| R_IPS | 14.96 | 5.44 | <.001 | **<.001** | <0.1 | -1.56 | -1.21 | .887 | ~1<sup>a</sup> | 12.12 | -2.08 | -1.93 | .973 | ~1 | 15.47 |

*Note:* The table shows the mean CCGP for each task component and ROI, the outputs (*Z* statistic, *p*-value) of the one-sided, one-sample Wilcoxon signed rank test addressing above-chance CCGP scores, and the adjusted *p-values* after a Bonferroni-Holm multiple comparisons correction. For each component and ROI, the table also includes the Bayes Factors supporting the null hypothesis (BF<sub>01</sub>) from one-sided one-sample Bayesian *t*-tests addressing above-chance CCGP scores (moderate to strong evidence for null effect in cursive).

In contrast, we did not find evidence supporting the presence of abstract neural codes for the target category or the relevant feature. Regarding the target category, the Bayesian T-tests provided moderate to strong evidence against the ROIs’ CCGP scores being greater than chance (see **Table 2**). The absence of pattern generalization was found systematically across all the cross-decoding iterations that led to the CCGP computation.

Finally, for the instructed relevant feature, Bayes Factors also provided moderate to strong support for the absence of coding generalization in most of the regions explored, except the R_aI/fO and the L_IFS, where this test was inconclusive. This effect was replicated across all the cross-decoding iterations, with a single exception, the L_IFS, where significant above-chance cross-decoding was found for only one of the context arrangements. Exploratory whole-brain searchlight analyses of CCGP showed a similar pattern of results. While a widespread cluster including areas within and outside the MDN exhibited significant CCGP for task demand (**Supp. Fig. 3**), no clusters survived the FWE correction for either target category or relevant feature (**Supp. Table 4**).

Last, and as a control analysis, we re-estimated the CCGP by embedding a cross-validation procedure within each cross-decoding protocol. With this approach, we sought to rule out that the CCGP scores were driven by overfitting of the classifiers to run-specific noise structure. These results broadly replicated the pattern of findings described above (**Supp. Table 3**). We found significant above-chance CCGP scores for the task demand in all the ROIs. Again, we did not detect any above-chance CCGP index for the target category nor the relevant feature component.

#### 3.2.3. Dimensionality of the encoding space

We estimated the dimensionality underlying the ROIs’ representational spaces by computing the SD, which measures the minimum number of significant above-chance classifications (here, interpreted as coding axes or dimensions) that can be performed in a dataset. This involved training binary classifiers to decode both informative and uninformative context dichotomies.

The results of this analysis (shown in Fig. 5A and **B**) suggested high-dimensional coding architectures across the MDN. In most regions, the spatial activity patterns were distributed not only following the instructions dimensions (those addressed in the decoding results) but also around non-interpretable axes that systematically separated different arrangements of task contexts. This manifested as statistically above-chance performance of classifiers decoding both task-informative and uninformative dichotomies (**Fig. 5A**). Despite this general finding, the shattering dimensionality also varied across MDN regions. It was numerically highest in the bilateral IPS and L_IFJ, and the pre-SMA/ACC, where we could recover 14 to 17 dimensions out of the 35 axes that could be estimated with this approach (L_IPS = 17, R_IPS = 15, L_IFJ = 17; pre-SMA/ACC = 14; **Fig. 5B**). The lowest dimensionality was in the left and right aI/fO, with estimated SD of 2 and 3, respectively.

**Figure 5.**
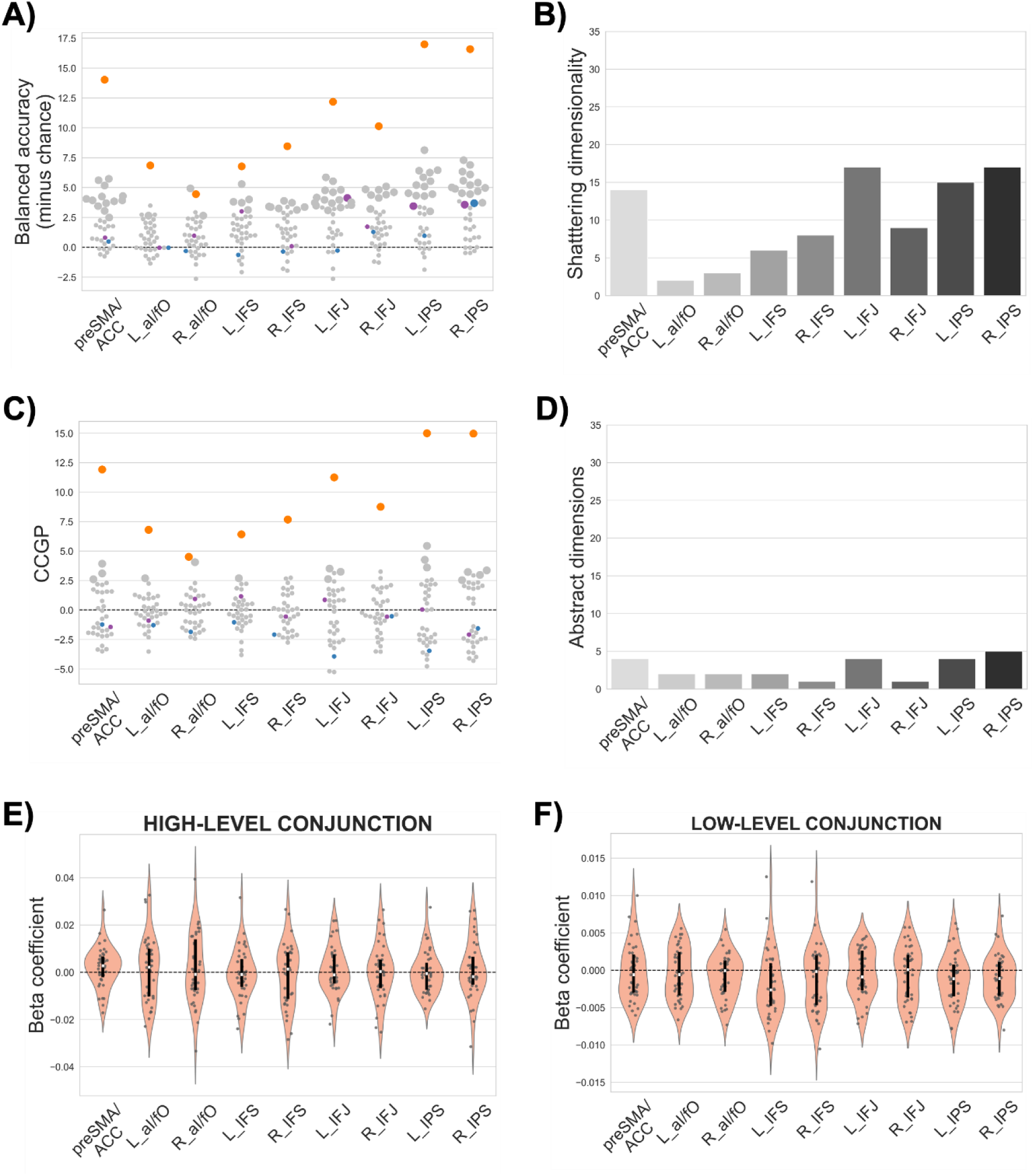
Coding geometry during novel task preparation. **(A, B)** Results from the Shattering Dimensionality (SD) analysis. The scatter plot **(A)** shows the balanced accuracies (minus chance) of the 35 dichotomies averaged across participants, with colored dots marking the three task components (task demand – orange, target category – blue, relevant feature – purple) and grey dots indicating task-irrelevant dimensions. Significantly above-chance decoding is shown with larger dots. The bar plot **(B)** depicts the estimated SD of each ROI, i.e., the count of significant classifications from **(A)**. **(C, D)** Results from the Cross-Condition Generalization Performance (CCGP) analysis applied to the 35 dichotomies. The scatter plot **(C)** shows the CCGP score of each dichotomy, and the bar plot **(D)** shows the count of dichotomies with significant CCGP scores per ROI, i.e., the number of abstractly coded dimensions. The plotting conventions are the same as in **(A, B). (E, F)** Results from the conjunctive models used in the Representational Similarity Analysis (RSA). Violin plots depict the ROIs’ beta coefficients for the high-level conjunctive model (**E**, combining task demand, target category and relevant feature) and the low-level conjunctive model obtained in a separate trial-wise RSA (**F,** combining specific feature and response mapping). Grey dots show individual participant means. The sample median is shown in the violins with a white dot. See **Supp.** Fig. 4 for the results of the remaining theoretical models tested.

Next, we aimed to further characterize these encoding axes by examining their level of abstractness and computed the CCGP scores for all the possible binary dichotomies (Fig. 5C and **D**). This analysis showed that many of the additional decodable dimensions found above did not generalise across task contexts. Instead, the main abstract and generalizable coding dimension was the task demand component. In most MDN ROIs, we detected either just the abstract task demand axis (R_IFS and R_IFJ) or a second, non-informative dimension (L_aI/fO, R_aI/fO, L_IFS). In the remaining regions, we detected four (R_IPS, L_IFJ, preSMA/ACC) or five (L_IPS) non-informative but abstract dimensions. Hence, despite the high dimensionality of MDN representational spaces, most of the non-informative coding axes were context-specific.

#### 3.2.4. Conjunctive task representations

Finally, we addressed the presence of conjunctive neural representations integrating the instructed task parameters into unique patterns. Following past studies (e.g., Kikumoto & Mayr, 2020), we employed model-based RSA to quantify how well the ROIs encoding spaces could be explained with theoretical models predicting pattern similarity only for same-context tasks, i.e., instructions sharing the same combination of the three task components.

After computing each ROI’s cross-validated RSM, we estimated multiple regressions using, as predictors, models based on the three task components and their conjunction. Results, shown in **Table 3**, replicated the overarching strong effect of task demand in shaping the encoding geometry, with this model linked to systematically above-zero beta weight in all the ROIs explored. Conversely, we did not detect a significant effect of the target category or the relevant feature with this analytical approach (see **Supp. Fig. 4A**). More critically, the conjunctive model did not explain task coding in any of the ROIs explored (see **Fig 5E**). Further Bayes Factors provided moderate to strong evidence supporting the null hypothesis in all ROIs except the preSMA/ACC, where inconclusive results were found (see **Table 3**). To ensure that the dominant effect of task demand was not obscuring more subtle effects of the other components or their conjunction, we conducted a follow-up exploratory analysis. We repeated the RSA as described in the Methods (Section 3.5.5), but separated the selection and integration conditions. This yielded isolated four-by-four RSMs, which were fitted with regressors for the target category, the relevant feature, and their conjunction. This control analysis mirrored our main findings: no significant effects emerged in any ROI for the main variables or the high-level conjunction (see **Supp. Fig. 5** and **Supp. Table 5**).

**Table 3.** Results from the model-based Representational Similarity Analysis (RSA) based on cross-validated Pearson correlation distances (from run-wise LSU beta maps).

|  | Task demands |  |  |  | Target category |  |  |  | Relevant feature |  |  |  | Conjunction |  |  |  | BF<sub>01</sub> |
| --- | --- | --- | --- | --- | --- | --- | --- | --- | --- | --- | --- | --- | --- | --- | --- | --- | --- |
|  | beta | z | p | p<sub>corr</sub> | beta | z | p | p<sub>corr</sub> | beta | z | p | p<sub>corr</sub> | beta | z | p | p<sub>corr</sub> |  |
| preSMA/ACC | 0.0086 | 4.85 | <.001 | <.001 | -0.0005 | -0.77 | .781 | ~1 | -0.0001 | -0.23 | .591 | ~1 | 0.0021 | 1.71 | .044 | .393 | 1.02 |
| L_al/fo | 0.0063 | 3.51 | <.001 | <.001 | -0.0005 | -0.15 | .558 | ~1 | 0.0003 | -0.08 | .531 | ~1 | 0.0012 | 0.31 | .377 | ~1 | 3.53 |
| R_al/fo | 0.0047 | 2.39 | .008 | .008 | -0.0006 | -0.98 | .837 | ~1 | 0.0002 | 0.64 | .263 | ~1 | 0.0012 | 0.36 | .361 | ~1 | 5.73 |
| L_IFS | 0.0093 | 4.25 | <.001 | <.001 | 0.0011 | 0.82 | .207 | ~1 | 0.0024 | 1.44 | .074 | .669 | -0.0008 | -0.79 | .785 | ~1 | 7.85 |
| R_IFS | 0.0112 | 4.63 | <.001 | <.001 | 0.0016 | 0.83 | .203 | ~1 | 0.0014 | 0.68 | .249 | ~1 | -0.0008 | -0.47 | .680 | ~1 | 7.70 |
| L_IFJ | 0.0121 | 5.04 | <.001 | <.001 | 0.0001 | -0.17 | .569 | ~1 | 0.0019 | 1.15 | .125 | .999 | 0.0009 | 0.30 | .382 | ~1 | 3.53 |
| R_IFJ | 0.0122 | 4.86 | <.001 | <.001 | 0.0021 | 2.03 | .021 | .190 | 0.0005 | 0.65 | .258 | ~1 | 0.0003 | 0.20 | .420 | ~1 | 5.09 |
| L_IPS | 0.0141 | 5.25 | <.001 | <.001 | 0.0005 | 0.09 | .464 | ~1 | 0.0013 | 0.83 | .203 | ~1 | 0 | -0.45 | .675 | ~1 | 5.75 |
| R_IPS | 0.0177 | 5.28 | <.001 | <.001 | 0.0028 | 1.63 | .052 | .416 | 0.0001 | 0.19 | .425 | ~1 | 0.0007 | 0.02 | .492 | ~1 | 4.27 |

*Note:* The table shows the mean beta weights for each RSA model and ROI, the outputs (Z statistic, *p*-value) of the one-sided, one-sample Wilcoxon signed rank test addressing above-zero regression coefficients, and the adjusted *p-values* after a Bonferroni-Holm multiple comparisons correction. For the conjunctive model, the table also includes the Bayes Factors supporting the null hypothesis (BF<sub>01</sub>) from one-sided one-sample Bayesian *t*-tests addressing above-zero beta weights (moderate to strong evidence for null effect in cursive).

The above analysis targeted higher-order task conjunctions, combining abstract components such as the current demands, supra-order categories, and features. In opposition, past studies reporting conjunctive coding entailed combinations between more concrete stimulus and response-oriented parameters. To provide more comparable results, we conducted an exploratory test targeting lower-level conjunctions conformed by the specific frame feature and response sets on each trial. For that, we estimated trial-by-trial activity patterns using an LSS method (see section 2.5) and estimated non-cross-validated, instruction-wise RSMs. The multiple regressions were fitted using, as predictors, models for the task demand, target category, relevant feature, specific feature, response set, and the low-level conjunction between the specific feature and the response set. These results (**Table 4**) replicated the above RSA regarding the task demand (with a significant effect in all ROIs) and the target category and relevant feature (neither predicting neural RSMs). Interestingly, both lower-level task parameters significantly explained the encoding geometry in several regions (see **Supp. Fig. 4B**). While the model conveying the specific frame color or feature was significant in the left aI/fO, IFJ, and IPS, the one coding the response mapping was significant in the IPS, bilaterally. Notwithstanding, the model predicting lower-level conjunctions, defined by the latter two task parameters, was not significant in any of the ROIs (see **Fig. 5F**). Bayesian one-sample *T*-tests corroborated that no low-level conjunction beta weight was greater than zero (**Table 4**).

**Table 4.** Results from the model-based Representational Similarity Analysis (RSA) based on trial-wise non-cross-validated Pearson correlation distances (from LSS beta maps).

|  | Task demands |  |  | Target category |  |  | Relevant feature |  |  | Specific feature |  |  | Resp. Mapping |  |  | Low-order conj. |  |  | BF<sub>01</sub> |
| --- | --- | --- | --- | --- | --- | --- | --- | --- | --- | --- | --- | --- | --- | --- | --- | --- | --- | --- | --- |
|  | z | p | p<sub>corr</sub> | z | p | p<sub>corr</sub> | z | p | p<sub>corr</sub> | z | p | p<sub>corr</sub> | z | p | p<sub>corr</sub> | z | p | p<sub>corr</sub> |  |
| preSMA/ACC | 5.18 | <.001 | <.001 | 0.91 | .180 | .541 | 0.24 | .404 | ~1 | 0.55 | .291 | .292 | 2.13 | .017 | .100 | -0.73 | .768 | ~1<sup>a</sup> | 7.59 |
| L_al/fO | 4.29 | <.001 | <.001 | 0.90 | .188 | .541 | -0.90 | .816 | ~1 | 2.74 | .003 | .024 | -0.02 | .508 | .508 | -0.82 | .793 | ~1<sup>a</sup> | 9.68 |
| R_al/fO | 3.90 | <.001 | <.001 | 0.75 | .228 | .377 | -1.13 | .872 | ~1 | 2.37 | .009 | .054 | 0.79 | .215 | .430 | -1.58 | .943 | ~1<sup>a</sup> | 15.15 |
| L_IFS | 5.30 | <.001 | <.001 | 1.60 | .056 | .275 | 1.14 | .128 | ~1 | 2.10 | .018 | .060 | 1.46 | .072 | .217 | -2.95 | .998 | ~1<sup>a</sup> | 19.98 |
| R_IFS | 5.00 | <.001 | <.001 | 1.14 | .128 | 0.510 | 0.47 | .320 | ~1 | 1.05 | .146 | .292 | 1.68 | .046 | .185 | -1.74 | .959 | ~1<sup>a</sup> | 14.64 |
| L_IFJ | 5.41 | <.001 | <.001 | 2.39 | .010 | .098 | 0.93 | .177 | ~1 | 2.90 | .002 | .017 | 1.90 | .028 | .142 | -0.57 | .719 | ~1<sup>a</sup> | 10.58 |
| R_IFJ | 5.38 | <.001 | <.001 | 2.28 | .012 | .097 | -1.15 | .875 | ~1 | 2.31 | .010 | .054 | 2.21 | .013 | .094 | -0.75 | .772 | ~1<sup>a</sup> | 9.98 |
| L_IPS | 5.44 | <.001 | <.001 | 2.06 | .020 | .138 | 0.80 | .211 | ~1 | 2.56 | .005 | .033 | 4.11 | <.001 | <.001 | -2.11 | .983 | ~1<sup>a</sup> | 16.73 |
| R_IPS | 5.44 | <.001 | <.001 | 1.84 | .033 | .199 | -0.67 | .751 | ~1 | 2.17 | .015 | .060 | 4.05 | <.001 | <.001 | -2.00 | .977 | ~1<sup>a</sup> | 15.94 |

*Note:* The table shows, for each RSA model and ROI, the outputs (Z statistic, *p*-value) of the one-sided, one-sample Wilcoxon signed rank test addressing above-zero regression coefficients, and the adjusted *p*-values after a Bonferroni-Holm multiple comparisons correction. For the conjunctive model, the table also includes the Bayes Factors supporting the null hypothesis (BF<sub>01</sub>) from one-sided one-sample Bayesian *t*-tests addressing above-zero beta weights (moderate to strong evidence for null effect in cursive).

## 4. Discussion

In this work, we aimed to characterize the neural coding mechanisms supporting novel task assembly across the MDN. To do so, we collected fMRI data while participants followed new verbal instructions and applied MVPA to address the informational content and format reflected in MDN activity. Addressing our first question, we identified distinct coding profiles across the network depending on the type of information manipulated: the overarching task demand, the relevant target category, and the visual features participants had to respond to. Next, focusing on the instruction-driven representational spaces, generalization and dimensionality analysis revealed both abstract and high-dimensional neural codes. Finally, we examined whether conjunctive instruction codes were present in MDN activity. However, model-based RSA did not provide evidence of such context-specific coding architecture. Overall, our findings suggest that combinatorial coding geometries within the MDN may play a role in novel, adaptive behavior, potentially exploiting both task generalization and expressivity, challenging the notion of an inherent trade-off between them.

In line with past research, our findings highlight that MDN activity is adaptively configured to encode task-relevant information (for a review, see Zheng, Lu & Woolgar, 2024 & Duncan, 2025). That is also the case in novel contexts, where these parameters are extracted from complex verbal commands (Cole et al., 2011; González-García et al., 2017, 2021; Palenciano et al., 2019a; Sobrado et al., 2022). The analyses showed different coding profiles within this network, depending on the specific task content investigated. On one hand, the overarching task demand component, indicating current selection vs. integration goals, was widely represented across the MDN. On the other hand, the encoding of the two stimulus-oriented parameters, the relevant target category and feature, was restricted to the IFJ and the IPS. Although this pattern is consistent with a functional specialization of these regions, their larger size (i.e., greater number of voxels) may also have increased decoding sensitivity, potentially contributing to the higher decoding accuracy observed. These findings are consistent with the idea that the MDN informational profile reflected the hierarchical task structure, with higher-level parameters implemented through widespread neural codes, and lower-level information being coded more locally. Supporting this view, recent EEG data obtained with the same paradigm revealed complementary temporal dynamics: while task demand was associated with temporally sustained and stable activity, target category and relevant feature were coded only transiently (Pena et al., 2025). More generally, similar hierarchical accounts have been formulated in the field of cognitive control (Badre, 2008, 2024; Koechlin et al., 2003), supported by both fMRI (Badre & Nee, 2018; Nee, 2021; Nee & D’Esposito, 2016) and electrophysiological (Cellier et al., 2022) evidence. Moreover, and resonating with the dual MDN model (Dosenbach et al., 2007, 2008), our findings highlighted that the lateral portion of this network (in comparison with the more medial, or cingulo-opercular one) conveyed more diverse information (Crittenden et al., 2016) from different hierarchical levels. While early research pointed towards the lateral prefrontal cortex as the key hub for representing multiplexed task information (Duncan, 2001; Miller & Cohen, 2001; Sakai, 2008), our work showed a similar scenario for the IPS, where the three task parameters were jointly encoded, in line with more recent network-based conceptualisations of cognitive control (Duncan et al., 2020; Cole et al., 2016; Karimi-Rouzbahani et al., 2026). Future research with paradigms spanning wider hierarchical information may be needed to further describe the coding profiles displayed across different parts of the MDN.

Our main goal was to address the geometry or structure governing novel task coding within the MDN by focusing on two core features identified in the literature: abstraction and dimensionality. The CCGP scores robustly evidenced coding abstraction, particularly for the task demand component, whose activity patterns systematically generalized across all possible context combinations throughout the MDN. This encoding format has been interpreted as the main signature of compositional coding, i.e., the recombination of common neural representations to build broader task models (Cole, Laurent, et al., 2013; Frankland & Greene, 2020; Reverberi et al., 2012; Tafazoli et al., 2026). Hence, it has been proposed as a key mechanism for novel instructed performance (Cole, Laurent, et al., 2013; Ito et al., 2022), with recent data showing its involvement during the first trials with a task (Mill et al., 2025). Through an exhaustive cross-decoding protocol, our work provided compelling evidence supporting this proposal, showing that patterns coding higher-order task information could be compositionally reused across new instructions. However, the same was not true for codes representing the lower-level target category or the relevant feature components. Although the two variables could be significantly decoded from IPS and IFJ activity patterns with the standard classification approach, we did not detect significant CCGP scores for either of them, nor above-chance accuracies in the individual cross-decoding iterations (only one significant result across 16 iterations and 9 ROIs, **Fig. 3C**). Interestingly, despite their transient nature, these lower-level task features showed evidence of abstraction in an EEG study with the same paradigm, albeit with a weaker effect than task demands (Pena et al., 2025). One possible explanation for these contrasting results is that the lower baseline decoding performance for these variables may have inherently reduced our statistical power to detect significant generalization effects. Additionally, transient task components may be difficult to detect with fMRI because of its relatively low temporal resolution. While this null finding should be treated with caution, similar results where generalizable and non-generalizable task dimensions coexist have been found in past studies (e.g., Sobrado et al., 2022). These findings suggest that compositional coding could be selectively recruited by specific sources of task content or contextual demands.

Critically, by examining both coding abstraction and dimensionality, we sought to understand the interplay between two possible geometries that could scaffold novel instruction following: either abstract and compressed (i.e., low-dimensional) architectures promoting generalization to novel contexts, or context-specific and high-dimensional ones, linked to greater read-out expressivity (Badre et al., 2021; Musslick & Cohen, 2021). Our data suggested a more complex picture, one in which abstract and generalizable neural codes coexisted alongside expansive, high-dimensional representational spaces. We predicted that neural code abstraction observed for overall task demand would be reflected in a compressed coding architecture. On the contrary, SD results showed that the distributed activity in the MDN was systematically structured around a greater number of axes than those required by the task, incorporating multiple arbitrary (or non-task-informative) dimensions. Numerically, this effect was rather generalized across the MDN. It is worth noting that, because the SD method provided a single dimensionality value per ROI (and not subject-wise estimates), these scores could not be statistically compared; thus, this finding should be interpreted with caution. Moreover, these high-dimensional estimates should be interpreted in light of the broad pattern of task demands in our data, which may have particularly affected some of the arbitrary dimensions. Nonetheless, such high SD estimates resonate with a wide body of literature showing expansive dimensionality in prefrontal and/or parietal activity patterns (Fusi et al., 2016), as has been shown in intracranial recordings (Bernardi et al., 2020; Rigotti et al., 2013) and fMRI (Bhandari et al., 2024). It has been thus considered an intrinsic property of these regions that maximizes the readout of their activity patterns and, hence, their expressivity (i.e., the number of distinguishable pieces of information that are encoded; Badre et al., 2021; Fusi et al., 2016; Rigotti et al., 2013). Furthermore, recent EEG results have suggested that encoding dimensionality is progressively expanded over the time course of task preparation to implement new, complex instructions (Pena et al., 2025). Taken together, novel task scenarios could recruit this geometrical feature across the MDN, leveraging greater dimensionalities to host diverse repertoires of instructed demands. Finally, in most ROIs, a subset of these arbitrary coding dimensions also displayed an abstract, generalizable format (**Fig. 4D**). Previous theoretical proposals have posited that abstract codes should be low dimensional (Badre et al., 2021; Musslick & Cohen, 2021), as a lower-dimensional structure would offer the most parsimonious solution for generalization. Our results, however, challenge this account by raising the possibility that more demanding coding organizations may also support abstraction. Consistent with this view, recent evidence suggests that generalization can emerge under associative inference learning, where latent task reinstatement has been observed (Lee et al, 2026). Although the interpretation remains uncertain, converging evidence from earlier work (Bernardi et al., 2020), points to a promising path for future research on coding geometry.

Our third question was whether conjunctive codes driven by novel instructions manifested in our paradigm, since recent studies have stressed the relevance of this coding architecture for adaptive task control (Kikumoto et al., 2024; Kikumoto, Mayr, et al., 2022; Kikumoto, Sameshima, et al., 2022; Kikumoto & Mayr, 2020; Chai et al., 2026; Lee et al., 2026). Conjunctive coding states that different task components are bound or mixed into unique representations (Hommel, 2009). This mechanism is tightly linked with high-dimensional spaces, where the dimensions are thought to implement these context-unique and segregated neural task codes (Badre et al., 2021). Nonetheless, we did not detect evidence supporting conjunctive novel task representations. In this regard, although we employed the standard approach used in the past to identify the presence of conjunctive coding at the macroscopic scale (model-based RSA and a similar theoretical RSM; e.g., Kikumoto & Mayr, 2020), our paradigm imposed a key difference. Specifically, previous studies have employed simpler tasks where a limited set of individual conjunctions (or task combinations) were widely repeated. The complexity of our paradigm, instead, led to different levels of conjunctions: from broader combinations of the three main task parameters to lower-level associations between specific features and response sets, but also finer-grained trial-wise combinations of instructed content. Our methodology allowed us to interrogate the two former possibilities (high- and low-level conjunction), not finding significant results for any. Still, we could not address whether individual instructions (shown only once in our paradigm) were conjunctively encoded into unique activity patterns. Moreover, past studies supporting conjunctive neural codes relied on temporally resolved EEG recordings. In contrast, the fMRI BOLD signal collected in our work integrates neural events at a substantially slower timescale, which could have hindered the detection of this phenomenon. Therefore, it remains uncertain whether our null results indicate a lack of conjunctive process for novel tasks or, instead, whether our design was not sensitive to the task content or the temporal scale at which these conjunctions are formed.

Several limitations of the current study should be acknowledged. Clearly, the slow timescale of fMRI BOLD signal constitutes a general constraint, as this technique is largely blind to fast changes in representational dynamics. Nevertheless, a previous EEG study using the same paradigm replicates several of our main findings, particularly the stability of higher-level task demands relative to the more transient effects of stimulus-bound variables (Pena et al., 2025). Related to the fMRI temporal precision, a further limitation concerns the proximity between epoch events: our design does not allow for a full dissociation between instruction, preparation and execution phases, leaving some uncertainty about the precise time windows in which these effects emerge. However, given that our primary analyses were not focused on contrasting these events, we do not consider this a critical issue (see Palenciano et al., 2019 or Muhle-Karbe et al., 2017 for a similar event design). Lastly, exploratory whole-brain analysis of information content (**Supp. Fig. 2** and **Supp. Table 2**) and generalization (**Sup. Fig. 3** and **Supp. Table 4**) revealed a widespread effect of task demand across the brain, alongside very small or null results for target category and relevant feature. This unexpected pattern may point to a potential bias toward task demand in our paradigm that could have overshadowed lower-level variables, although we implemented multiple controls during both task design and analysis to prevent potential confounds. To test whether effects for lower-level variables might emerge when accounting for task demand, we exploratorily performed RSA separately within each task demand condition. The null results persisted for target category, relevant feature, and their conjunction across all ROIs (see **Supp. Fig. 5** and **Supp. Table 5**). This control analysis therefore, does not provide evidence that task demand confounding explains the observed pattern. To further assess whether task demand effects reflected overall task difficulty (i.e., more challenging in selection; see **Fig. 3**), we correlated subject-wise behavioral measures (accuracy and reaction times) with mean task demand decoding across ROIs. This analysis again revealed no significant effects after multiple-comparison correction, and Bayes factors provided moderate to strong evidence for the null in most ROIs (see **Supp. Table 6**). Notably, this widespread task effect aligns with recent accounts emphasizing distributed neural effects during complex behavior (International Brain Laboratory, 2025; Mill et al., 2025), suggesting that the difficulty imposed by our current paradigm may underlie such broad representations of task demand. Moreover, EEG results with the same paradigm also corroborate strong and stable task coding (Pena et al., 2025).

Overall, our findings suggest a complex interplay of representational features underpinning novel task assembly. Although most of the current theoretical models state that the two coding geometries we explored (abstract and compact vs. context-unique and high-dimensional) constitute the two extremes of a continuum (Badre et al., 2021; Musslick & Cohen, 2021; but also see Egner, 2023), recent studies have also reported the presence of blended architectures that combine abstraction with high-dimensionality (Bernardi et al., 2020; Pena et al., 2025). Hence, independently combining both coding principles could effectively tailor the coding geometry to current challenges, achieving a balance between generalization and expressivity. In our case, the abstract representational format found in MDN areas for task demand may facilitate the transfer of acquired task representations to novel contexts, whereas using high dimensionality to encode novel rules could also account for the expressivity that is inherent to flexible, novel behavior. Notably, although the presence of abstract neural codes and high-dimensional spaces was rather ubiquitous across the MDN, we found a trend towards numerically greater estimates converging in the IPS and IFJ (i.e., the regions displaying more diverse informational content) and preSMA/ACC. While our current data do not allow us to extract further conclusions about MDN dynamics, it opens the possibility that an in-depth characterization of the representational geometry could advance our mechanistic understanding of this brain network.

## Data and Code Availability

Task and analysis scripts will be made available upon publication. Raw data will be uploaded at publication.

## Acknowledgements

This research was supported by grant PID2022-138940NB-100 awarded to MR and grant PID2023-151911NA-I00 awarded to AFP, funded by MCIN/EI/10.13039/501100011033/ and FEDER, UE. AFP was supported by Grant PAIDI21_00207 of the Andalusian Autonomic Government. AW was supported by intramural funding from the UK Research and Innovation Medical Research Council, grant number MC_UU_00030/15. CCG was supported by Project PID2023-149428NB-I00 funded by MICIU/AEI/10.13039/501100011033 and by ERDF/EU, and Grant RYC2021-033536-I funded by MCIN/AEI/10.13039/501100011033 and by the European Union NextGeneration EU/PRTR. PP was supported by scholarship PREP2023-002013, funded by MCIU/AEI/10.13039/501100011033 and FSE+. The Mind, Brain and Behavior Research Center receives funding from grants CEX2023-001312-M by MCIN/AEI/10.13039/501100011033 and UCE-PP2023-11 by the University of Granada. We are grateful to Marta Becerra Losada for assistance with data collection.

## References

Arco, J. E., González-García, C., Díaz-Gutiérrez, P., Ramírez, J., & Ruz, M. (2018). Influence of activation pattern estimates and statistical significance tests in fMRI decoding analysis. Journal of Neuroscience Methods, 308, 248–260. 10.1016/j.jneumeth.2018.06.017

Assem, M., Glasser, M. F., Van Essen, D. C., & Duncan, J. (2020). A domain-general cognitive core defined in multimodally parcellated human cortex. Cerebral Cortex, 30(8), 4361–4380. 10.1093/cercor/bhaa023

Badre, D. (2008). Cognitive control, hierarchy, and the rostro-caudal organization of the frontal lobes. In Trends in Cognitive Sciences (Vol. 12, Issue 5, pp. 193–200). 10.1016/j.tics.2008.02.004

Badre, D. (2024). Cognitive Control. Annual Review of Psychology Downloaded from Www.Annualreviews.Org. University of Gent, 12(6), 34. 10.1146/annurev-psych-022024

Badre, D., Bhandari, A., Keglovits, H., & Kikumoto, A. (2021). The dimensionality of neural representations for control. In Current Opinion in Behavioral Sciences (Vol. 38, pp. 20–28). Elsevier Ltd. 10.1016/j.cobeha.2020.07.002

Badre, D., & Nee, D. E. (2018). Frontal Cortex and the Hierarchical Control of Behavior. 10.1016/j.tics.2017.11.005

Bernardi, S., Benna, M. K., Rigotti, M., Munuera, J., Fusi, S., & Salzman, C. D. (2020). The Geometry of Abstraction in the Hippocampus and Prefrontal Cortex. Cell, 183(4), 954–967.e21. 10.1016/j.cell.2020.09.031

Bhandari, A., Keglovits, H., & Badre, D. (2024). Task structure tailors the geometry of neural representations in human lateral prefrontal cortex. BioRxiv, 2024.03.06.583429. 10.1101/2024.03.06.583429

Brass, M., Liefooghe, B., Braem, S., & De Houwer, J. (2017). Following new task instructions: Evidence for a dissociation between knowing and doing. Neuroscience & Biobehavioral Reviews, 81, 16–28. https://linkinghub.elsevier.com/retrieve/pii/S0149763416306856

Brincat, S. L., Siegel, M., von Nicolai, C., & Miller, E. K. (2018). Gradual progression from sensory to task-related processing in cerebral cortex. Proceedings of the National Academy of Sciences of the United States of America, 115(30), E7202–E7211. 10.1073/PNAS.1717075115/SUPPL_FILE/PNAS.1717075115.SAPP.P DF

Cellier, D., Petersen, I. T., & Hwang, K. (2022). Dynamics of Hierarchical Task Representations. 10.1523/JNEUROSCI.0233-22.2022

Chai, M., Ikink, I., Mattioni, S., Benavides, R. A., Kukkonen, N., Senoussi, M., … & Braem, S. (2026). Parallel control of conjunctive and compositional representations supports dynamic task preparation. bioRxiv, 2026-01. 10.64898/2026.01.26.701713

Cole, M. W., Bagic, A., Kass, R., & Schneider, W. (2010). Prefrontal dynamics underlying rapid instructed task learning reverse with practice. Journal of Neuroscience, 30(42), 14245–14254. 10.1523/JNEUROSCI.1662-10.2010

Cole, M. W., Etzel, J. A., Zacks, J. M., Schneider, W., & Braver, T. S. (2011). Rapid Transfer of Abstract Rules to Novel Contexts in Human Lateral Prefrontal Cortex. Frontiers in Human Neuroscience, 5(November), 1–13. 10.3389/fnhum.2011.00142

Cole, M. W., Ito, T., & Braver, T. S. (2016). The Behavioral Relevance of Task Information in Human Prefrontal Cortex. Cerebral Cortex, 26(6), 2497–2505. 10.1093/cercor/bhv072

Cole, M. W., Laurent, P., & Stocco, A. (2013). Rapid instructed task learning: A new window into the human brain’s unique capacity for flexible cognitive control. *Cognitive*, Affective and Behavioral Neuroscience, 13(1), 1–22. 10.3758/s13415-012-0125-7

Cole, M. W., Reynolds, J. R., Power, J. D., Repovs, G., Anticevic, A., & Braver, T. S. (2013). Multi-task connectivity reveals flexible hubs for adaptive task control. Nature Neuroscience, 16(9), 1348–1355. 10.1038/nn.3470

Crittenden, B. M., Mitchell, D. J., & Duncan, J. (2016). Task Encoding across the Multiple Demand Cortex Is Consistent with a Frontoparietal and Cingulo-Opercular Dual Networks Distinction. The Journal of Neuroscience: The Official Journal of the Society for Neuroscience, 36(23), 6147–6155. 10.1523/JNEUROSCI.4590-15.2016

Dosenbach, N. U. F., Fair, D. A., Cohen, A. L., Schlaggar, B. L., & Petersen, S. E. (2008). A dual-networks architecture of top-down control. Trends in Cognitive Sciences, 12(3), 99–105. https://www.ncbi.nlm.nih.gov/pmc/articles/PMC3632449/pdf/nihms-452910.pdf

Dosenbach, N. U. F., Fair, D. A., Miezin, F. M., Cohen, A. L., Wenger, K. K., Dosenbach, R. A. T., Fox, M. D., Snyder, A. Z., Vincent, J. L., Raichle, M. E., Schlaggar, B. L., & Petersen, S. E. (2007). Distinct brain networks for adaptive and stable task control in humans. Proceedings of the National Academy of Sciences, 104(26), 11073–11078. 10.1073/pnas.0704320104

Duncan, J. (2001). An adaptive coding model of neural function in prefrontal cortex. Nature Reviews Neuroscience, 2(11), 820–829. 10.1038/35097575

Duncan, J. (2010). The multiple-demand (MD) system of the primate brain: mental programs for intelligent behaviour. Trends in Cognitive Sciences, 14(4), 172–179. 10.1016/j.tics.2010.01.004

Duncan, J., Assem, M., & Shashidhara, S. (2020). Integrated intelligence from distributed brain activity. Trends in cognitive sciences, 24(10), 838–852. 10.1016/j.tics.2020.06.012

Duncan, J. (2025). Construction and use of mental models: Organizing principles for the science of brain and mind. Neuropsychologia, 207, 109062. 10.1016/j.neuropsychologia.2024.109062

Egner, T. (2023). Principles of cognitive control over task focus and task switching. Nature Reviews Psychology, 2(11), 702–714. 10.1038/s44159-023-00234-4

Fedorenko, E., Duncan, J., & Kanwisher, N. (2013). Broad domain generality in focal regions of frontal and parietal cortex. Proceedings of the National Academy of Sciences of the United States of America, 110(41), 16616–16621. 10.1073/pnas.1315235110

Frankland, S. M., & Greene, J. D. (2020). Concepts and Compositionality: In Search of the Brain’s Language of Thought. Annual Review of Psychology, 71, 273–303. 10.1146/ANNUREV-PSYCH-122216-011829

Freund, M. C., Etzel, J. A., & Braver, T. S. (2021). Neural coding of cognitive control: The representational similarity analysis approach. Trends in Cognitive Sciences, 25(7), 622. 10.1016/J.TICS.2021.03.011

Fusi, S., Miller, E. K., Rigotti, M., Karpova, A., & Kiani, R. (2016). Why neurons mix: high dimensionality for higher cognition This review comes from a themed issue on Neurobiology of behavior. Current Opinion in Neurobiology, 37, 66–74. 10.1016/j.conb.2016.01.010

González-García, C., Arco, J. E., Palenciano, A. F., Ramírez, J., & Ruz, M. (2017). Encoding, preparation and implementation of novel complex verbal instructions. NeuroImage, 148, 264–273. 10.1016/J.NEUROIMAGE.2017.01.037

González-García, C., Formica, S., Wisniewski, D., & Brass, M. (2021). Frontoparietal action-oriented codes support novel instruction implementation. NeuroImage, 226, 117608. 10.1016/j.neuroimage.2020.117608

Gorgolewski, K. J., Auer, T., Calhoun, V. D., Craddock, R. C., Das, S., Duff, E. P., Flandin, G., Ghosh, S. S., Glatard, T., Halchenko, Y. O., Handwerker, D. A., Hanke, M., Keator, D., Li, X., Michael, Z., Maumet, C., Nichols, B. N., Nichols, T. E., Pellman, J., … Poldrack, R. A. (2016). The brain imaging data structure, a format for organizing and describing outputs of neuroimaging experiments. Scientific Data 2016 3:1, 3(1), 1–9. 10.1038/sdata.2016.44

Hartstra, E., Kühn, S., Verguts, T. and Brass, M. (2011). The implementation of verbal instructions: An fMRI study. Hum. Brain Mapp., 32, 1811–1824. 10.1002/hbm.21152

Haxby, J. V., Connolly, A. C., & Guntupalli, J. S. (2014). Decoding Neural Representational Spaces Using Multivariate Pattern Analysis. Annual Review of Neuroscience, 37(1), 435–456. 10.1146/annurev-neuro-062012-170325

Haynes, J. D., & Rees, G. (2006). Decoding mental states from brain activity in humans. In Nature Reviews Neuroscience (Vol. 7, Issue 7, pp. 523–534). Nature Publishing Group. 10.1038/nrn1931

Hebart, M. N., Görgen, K., & Haynes, J.-D. (2014). The Decoding Toolbox (TDT): a versatile software package for multivariate analyses of functional imaging data. Frontiers in Neuroinformatics, 8, 88. 10.3389/fninf.2014.00088

Holm, S. (1979). A Simple Sequentially Rejective Multiple Test Procedure. Scandinavian Journal of Statistics, 6(2), 65–70. https://www.jstor.org/stable/4615733?seq=1#metadata_info_tab_contents

Hommel, B. (2009). Action control according to TEC (theory of event coding). Psychological Research, 73(4), 512–526. 10.1007/s00426-009-0234-2

International Brain Laboratory., Angelaki, D., Benson, B. et al. A brain-wide map of neural activity during complex behaviour. Nature 645, 177–191 (2025). 10.1038/s41586-025-09235-0

Jackson, J. B., Rich, A. N., Moerel, D., Teichmann, L., Duncan, J., & Woolgar, A. (2025). Domain general frontoparietal regions show modality-dependent coding of auditory and visual rules. Imaging Neuroscience, 3, IMAG.a.29. 10.1162/IMAG.a.29

Karimi-Rouzbahani, H., Rich, A. N., & Woolgar, A. (2026). Spatiotemporal characterisation of information coding in the multiple demand network. Imaging Neuroscience, 4, IMAG-a. 10.1162/imag.a.1171

Kikumoto, A., Bhandari, A., Shibata, K. et al. A transient high-dimensional geometry affords stable conjunctive subspaces for efficient action selection. Nat Commun 15, 8513 (2024). 10.1038/s41467-024-52777-6

Kikumoto, A., & Mayr, U. (2020). Conjunctive representations that integrate stimuli, responses, and rules are critical for action selection. PNAS, 117(19), 10603–10608. 10.1073/pnas.1922166117/-/DCSupplemental

Kikumoto, A., Mayr, U., & Badre, D. (2022). The Role of Conjunctive Representations in Prioritizing and Selecting Planned Actions. ELife, 11, 1–44. 10.7554/ELIFE.80153

Kikumoto, A., Sameshima, T., & Mayr, U. (2022). The Role of Conjunctive Representations in Stopping Actions. Psychological Science, 33(2), 325–338. 10.1177/09567976211034505/ASSET/IMAGES/LARGE/10.1177_09567976211034505-FIG5.JPEG

Koechlin, E., Ody, C., & Kouneiher, F. (2003). The Architecture of Cognitive Control in the Human Prefrontal Cortex. Science, 302(5648), 1181–1185. 10.1126/SCIENCE.1088545/ASSET/67FEC198-C1DD-4C72-96E8-27A3AEF539CF/ASSETS/GRAPHIC/SE4532039005.JPEG

Kriegeskorte, N., & Kievit, R. A. (2013). Representational geometry: Integrating cognition, computation, and the brain. In Trends in Cognitive Sciences (Vol. 17, Issue 8, pp. 401–412). 10.1016/j.tics.2013.06.007

Lee, W. T., Hazeltine, E., & Jiang, J. (2026). Neural traces of composite tasks in complex task representation in the human brain reflects learning performance. PLoS biology, 24(1), e3003613.10.1371/journal.pbio.3003613

Mill, R. D., & Cole, M. W. (2025). Dynamically shifting from compositional to conjunctive brain representations supports cognitive task learning. Nature communications, 16(1), 10084. 10.1038/s41467-025-65041-2

Miller, E. K., & Cohen, J. D. (2001). An Integrative Theory of Prefrontal Cortex Function. Annual Review of Neuroscience, 24(1), 167–202. 10.1146/annurev.neuro.24.1.167

Misaki, M., Kim, Y., Bandettini, P. A., & Kriegeskorte, N. (2010). Comparison of multivariate classifiers and response normalizations for pattern-information fMRI. NeuroImage, 53(1), 103–118. 10.1016/J.NEUROIMAGE.2010.05.051

Muhle-Karbe, P. S., Duncan, J., De Baene, W., Mitchell, D. J., & Brass, M. (2017). Neural Coding for Instruction-Based Task Sets in Human Frontoparietal and Visual Cortex. Cerebral Cortex, 27(3), 1891–1905. 10.1093/cercor/bhw032

Mumford, J. A., Davis, T., & Poldrack, R. A. (2014). The impact of study design on pattern estimation for single-trial multivariate pattern analysis. NeuroImage, 103, 130–138. 10.1016/J.NEUROIMAGE.2014.09.026

Mumford, J. A., Turner, B. O., Ashby, F. G., & Poldrack, R. A. (2012). Deconvolving BOLD activation in event-related designs for multivoxel pattern classification analyses. NeuroImage, 59, 2636–2643. 10.1016/j.neuroimage.2011.08.076

Musslick, S., & Cohen, J. D. (2021). Rationalizing constraints on the capacity for cognitive control. In Trends in Cognitive Sciences (Vol. 25, Issue 9, pp. 757–775). Elsevier Ltd. 10.1016/j.tics.2021.06.001

Nee, D. E. (2021). Integrative frontal-parietal dynamics supporting cognitive control. ELife, 10. 10.7554/ELIFE.57244

Nee, D. E., & D’Esposito, M. (2016). The hierarchical organization of the lateral prefrontal cortex. ELife, 5. 10.7554/eLife.12112

Norman, D. A., & Shallice, T. (1986). Attention to Action. In R. J. Davidson, G. E. Schwartz, & D. Shapiro (Eds.), Consciousness and Self-Regulation: Advances in Research and Theory Volume 4 (pp. 1–18). Springer US. 10.1007/978-1-4757-0629-1_1

Palenciano, A. F., González-García, C., Arco, J. E., Pessoa, L., & Ruz, M. (2019a). Representational organization of novel task sets during proactive encoding. Journal of Neuroscience, 0725–19. 10.1523/JNEUROSCI.0725-19.2019

Palenciano, A. F., González-García, C., Arco, J. E., & Ruz, M. (2019b). Transient and sustained control mechanisms supporting novel instructed behavior. Cerebral Cortex, 29(9), 3948–3960. 10.1093/cercor/bhy273

Palenciano, A. F., González-García, C., De Houwer, J., Liefooghe, B., & Brass, M. (2024). Concurrent response and action effect representations across the somatomotor cortices during novel task preparation. Cortex, 177, 150–169. 10.1016/j.cortex.2024.05.003

Pena, P., Palenciano, A. F., González-García, C., & Ruz, M. (2025). Novel verbal instructions recruit abstract neural patterns of time-variable information dimensionality. Journal of Neuroscience, 45(17). 10.1523/JNEUROSCI.1964-24.2025

Penny, W. D., Friston, K. J., Ashburner, J. T., Kiebel, S. J., & Nichols, T. E. (Eds.). (2006). Statistical Parametric Mapping: the Analysis of Functional Brain Images. Academic Press [Imprint].

Reverberi, C., Görgen, K., & Haynes, J.-D. (2012). Compositionality of Rule Representations in Human Prefrontal Cortex. Cerebral Cortex, 22(6), 1237–1246. 10.1093/cercor/bhr200

Rigotti, M., Barak, O., Warden, M. R., Wang, X.-J., Daw, N. D., Miller, E. K., & Fusi, S. (2013). The importance of mixed selectivity in complex cognitive tasks. Nature, 497(7451), 585–590. 10.1038/nature12160

Ritchie, J. B., Lee Masson, H., Bracci, S., & Op de Beeck, H. P. (2021). The unreliable influence of multivariate noise normalization on the reliability of neural dissimilarity. NeuroImage, 245, 118686. 10.1016/J.NEUROIMAGE.2021.118686

Sakai, K. (2008). Task Set and Prefrontal Cortex. Annual Review of Neuroscience, 31(1), 219–245. 10.1146/annurev.neuro.31.060407.125642

Schmalz, X., Biurrun Manresa, J., & Zhang, L. (2023). What is a Bayes factor?. Psychological methods, 28(3), 705.

Shashidhara, S., Assem, M., Glasser, M. F., & Duncan, J. (2024). Task and stimulus coding in the multiple-demand network. Cerebral Cortex, 34(7), bhae278. 10.1093/cercor/bhae278

Sobrado, A., Palenciano, A. F., González-García, C., & Ruz, M. (2022). The effect of task demands on the neural patterns generated by novel instruction encoding. Cortex, 149, 59–72. 10.1016/J.CORTEX.2022.01.010

Tafazoli, S., Bouchacourt, F.M., Ardalan, A. et al. Building compositional tasks with shared neural subspaces. Nature 650, 164–172 (2026). 10.1038/s41586-025-09805-2

Walther, A., Nili, H., Ejaz, N., Alink, A., Kriegeskorte, N., & Diedrichsen, J. (2016). Reliability of dissimilarity measures for multi-voxel pattern analysis. NeuroImage, 137, 188–200. 10.1016/j.neuroimage.2015.12.012

Waskom, M. L. (2021). seaborn: statistical data visualization. Journal of Open Source Software, 6(60), 3021. 10.21105/JOSS.03021

Woolgar, A., Jackson, J., & Duncan, J. (2016). Coding of Visual, Auditory, Rule, and Response Information in the Brain: 10 Years of Multivoxel Pattern Analysis. Journal of Cognitive Neuroscience, 28(10), 1433–1454. 10.1162/jocn_a_00981

Woolgar, A., Williams, M. A., & Rich, A. N. (2015). Attention enhances multi-voxel representation of novel objects in frontal, parietal and visual cortices. Neuroimage, 109, 429–437. 10.1016/j.neuroimage.2014.12.083

Zheng, Y., Lu, R., & Woolgar, A. (2024). Radical flexibility of neural representation in frontoparietal cortex and the challenge of linking it to behaviour. Current Opinion in Behavioral Sciences, 57, 101392. 10.1016/j.cobeha.2024.101392

