## Supplementary_Materials for "Multiple-Demand Network encoding geometry balances generalization and dimensionality during novel task assembly"

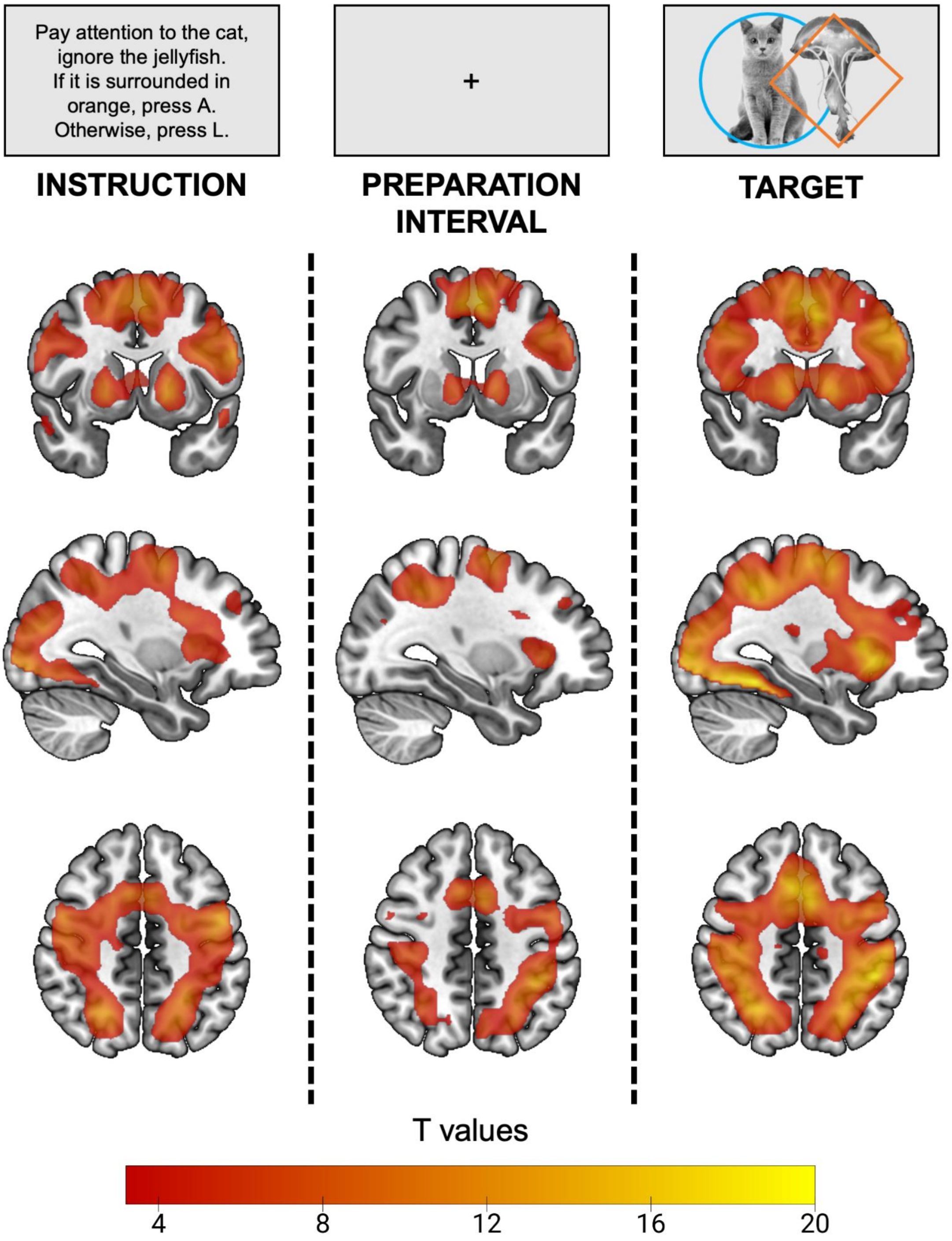


**S-Figure 1.** Univariate signal increase (relative to the implicit baseline) linked to the three events of the paradigm: instruction encoding (left column), preparation interval (middle column), and target presentation (right column). The figure shows cluster-wise FWE corrected (at $p < .05$, from an initial cluster-forming threshold of $p < .001$) whole brain T-maps. The color scale reflects the T values. Brain maps are shown at MNI coordinates [-31, 6, 46]. Unthresholded whole-brain maps are available on NeuroVault: https://neurovault.org/collections/JJORCZYL/

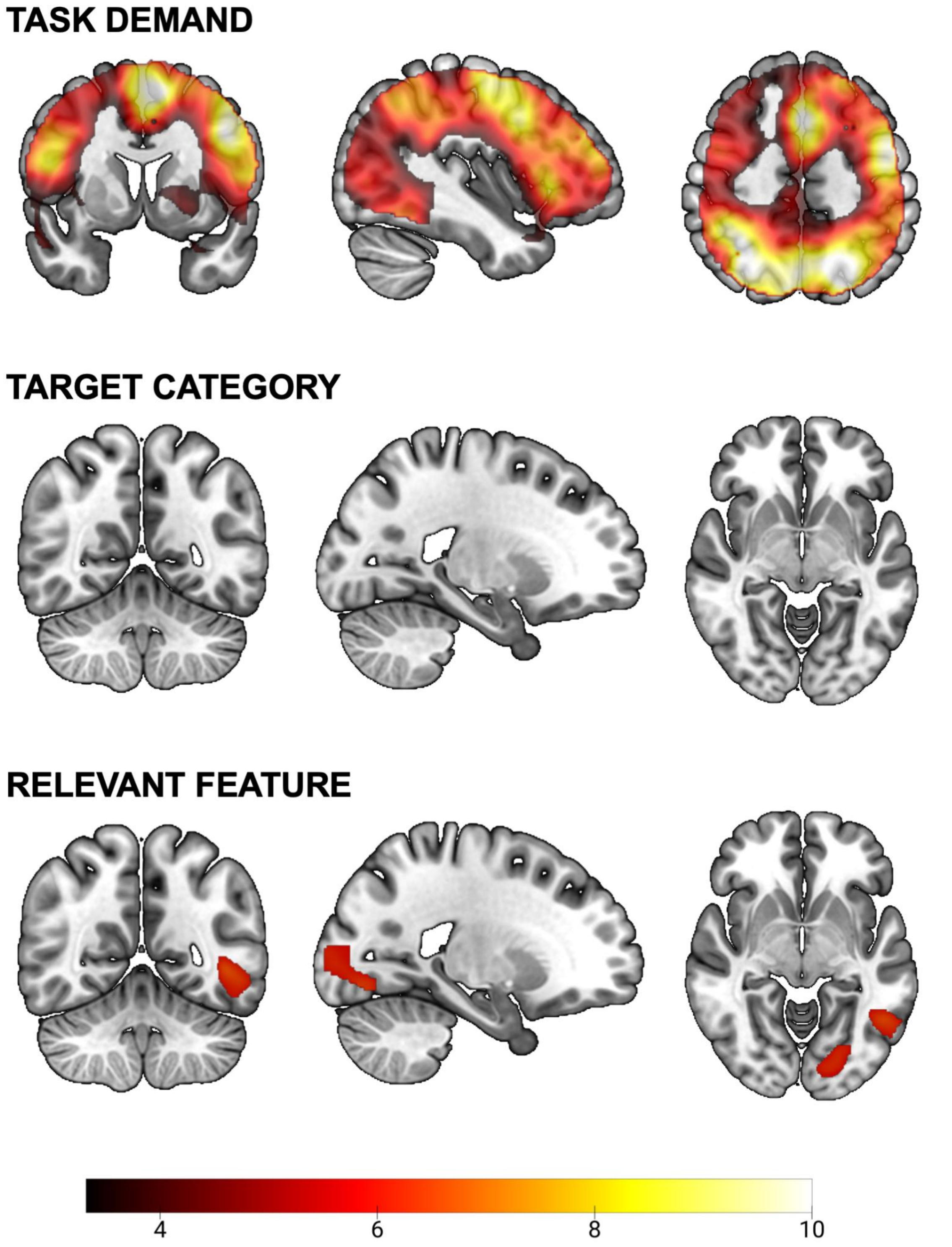


**S-Figure 2.** Searchlight whole-brain maps of neural decoding. Maps display whole-brain t-statistics thresholded using cluster-wise FWE correction at $p < .05$, based on an initial cluster-forming threshold of $p < .001$. Significant clusters of cross-validated balanced accuracy minus chance are shown for Task Demand (upper row; MNI coordinates: [-40, 6, 49]), Target Category (middle row; MNI coordinates: [-22, -55, -6]) and Target Relevant Feature (lower row; MNI coordinates: [-22, -55, -6]). No clusters survived correction for Target Category. Unthresholded whole-brain maps are available on NeuroVault: https://neurovault.org/collections/JJORCZYL/

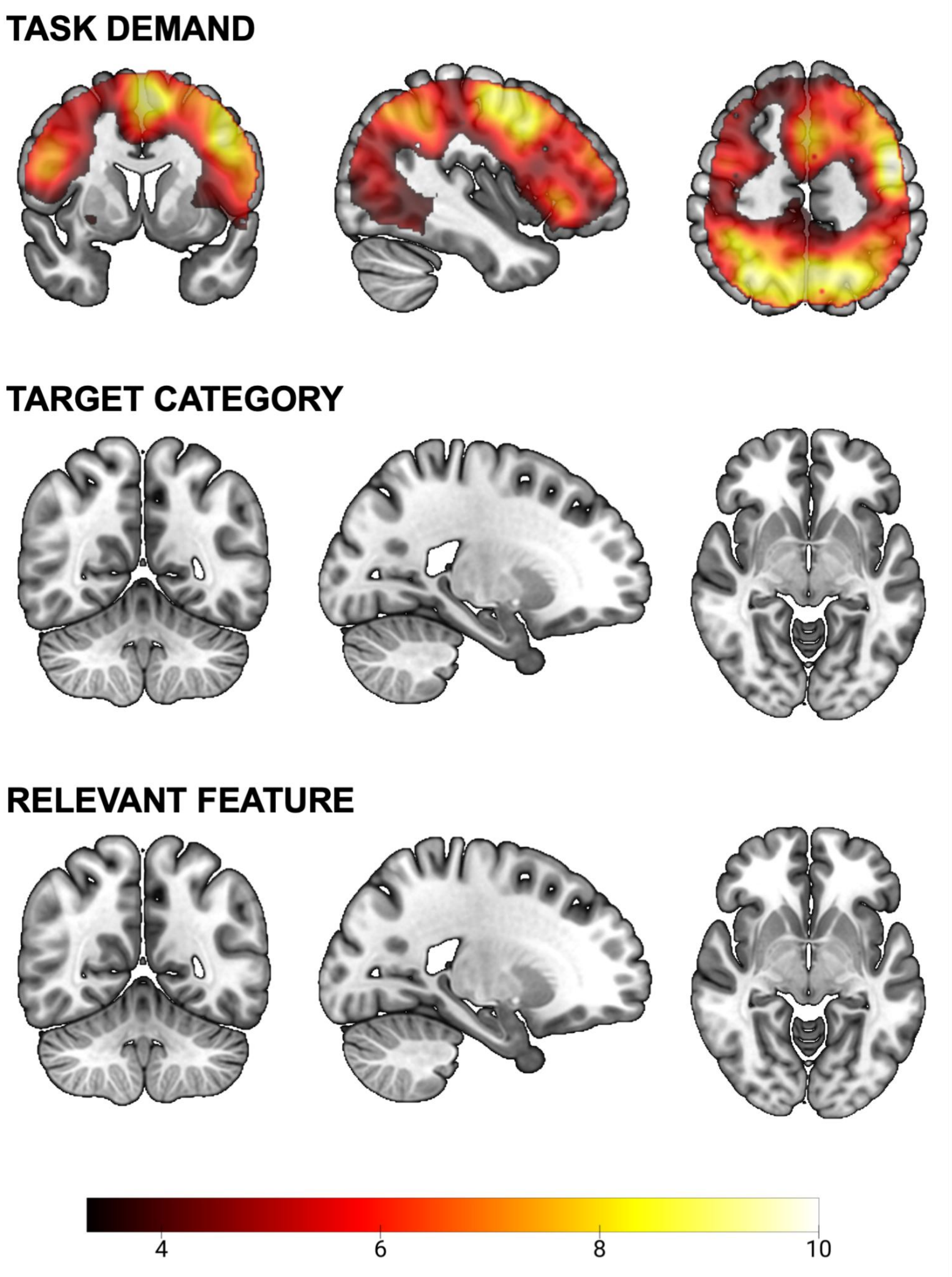


**S-Figure 3.** Searchlight whole-brain maps of Cross-Condition Generalization Performance (CCGP). Maps display whole-brain t-statistics thresholded using cluster-wise FWE correction at $p$ < .05, based on an initial cluster-forming threshold of $p$ < .001. Significant clusters of CCGP are shown for Task Demand (upper row; MNI coordinates: [-40, 6, 49]), Target Category (middle row; MNI coordinates: [-22, -55, -6]) and Target Relevant Feature (lower row; MNI coordinates: [-22, -55, -6]). No clusters survived correction for Target Category and Relevant Feature. Unthresholded whole-brain maps are available on NeuroVault: https://neurovault.org/collections/JJORCZYL/

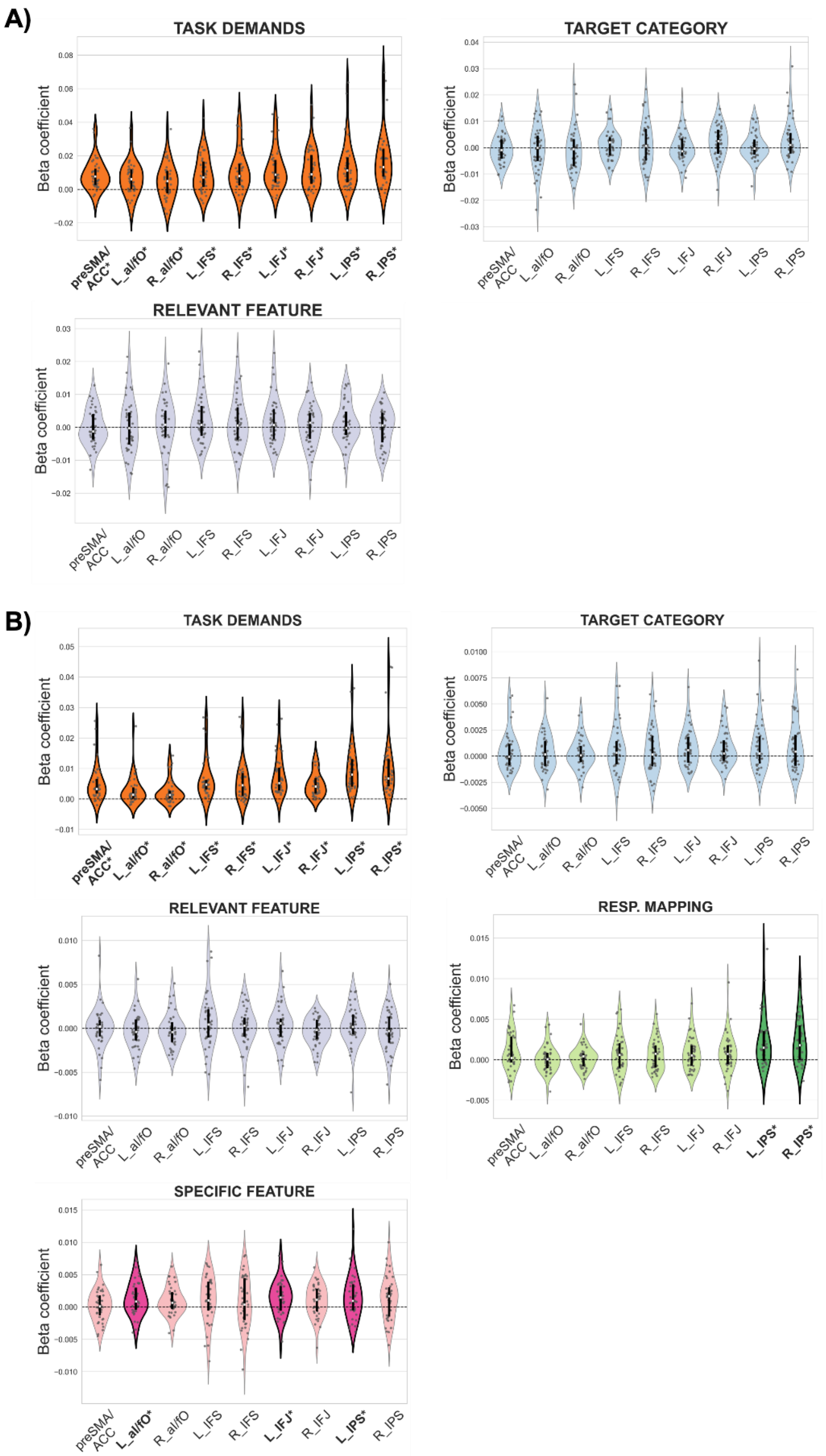


**S-Figure 4.** Results from Representational Similarity Analysis (RSA) complementary to the conjunctive models shown in Figure 5E-F. Violin plots depict the ROIs' beta coefficients for theoretical models tested alongside the high-level conjunction model in RSA (**A**, combining task demand, target category, and relevant feature) and the low-level conjunctive model derived from a separate trial-wise RSA (**B,** combining task demand, target category, relevant feature, specific feature, and response mapping). Grey dots represent individual participant means. The sample median is indicated by the white dot within each violin.

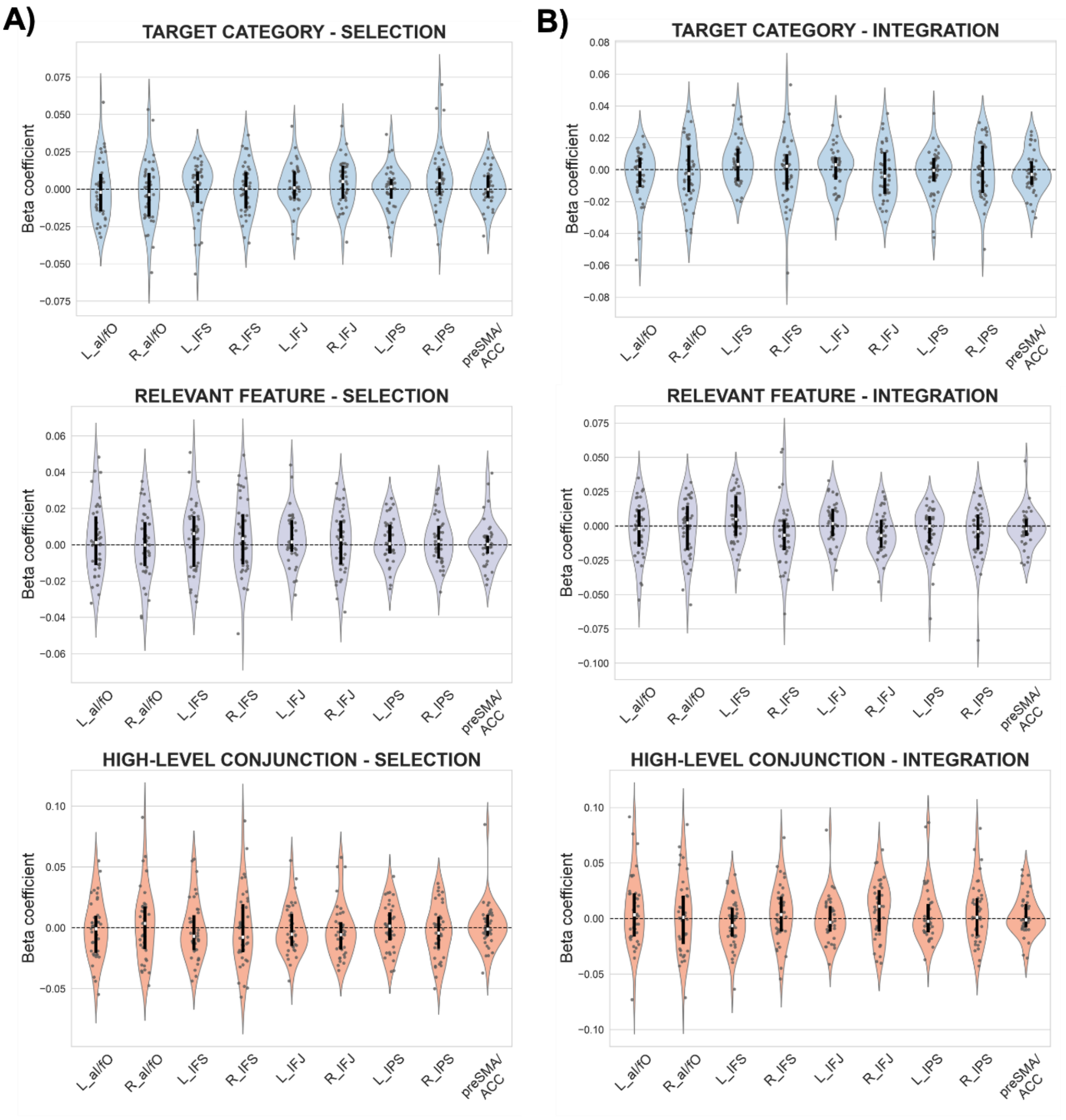


**S-Figure 5.** Results from Representational Similarity Analysis (RSA) separated by task demand (from run-wise LSU beta maps). This control analysis was conducted to ensure that the effects of task demand did not obscure subtle effects of other variables. We performed model-based RSA condition-wise as described in Section 2.5.5, separately for integration and selection conditions, using four-by-four representational similarity matrices. (**A, B**) Violin plots depict the ROIs' beta coefficients for the selection condition (**A**) and the integration condition (**B**). Grey dots indicate individual participant means. The sample median is shown in the violins with a white dot.

**Supplementary Table 1**.

Results from the univariate whole-brain, one-sample *t*-test contrasting each paradigm epoch regressor against the implicit baseline.

| Contrast | Regions | Hemisphere | X | Y | Z | k | *t* value |
|---|---|---|---|---|---|---|---|
| **Instruction** | MDN, ventral visual stream, motor and somatosensory networks, thalamus, basal ganglia (putamen, caudate, globus pallidus) | Bilateral | 26 | -96 | -4 | 47189 | 18.56 |
| | Posterior cingular cortex | Bilateral | 8 | -42 | 22 | 564 | 6.95 |
| | Superior, middle frontal gyrus | Right | 30 | 40 | 32 | 354 | 6.43 |
| | Superior, middle temporal gyri | Right | 48 | -26 | 2 | 1252 | 6.42 |
| | Ventromedial prefrontal cortex | Bilateral | 2 | 58 | -16 | 455 | 5.10 |
| **Preparation interval** | MDN, thalamus, basal ganglia (putamen, caudate, globus pallidus) | Bilateral | -4 | 6 | 52 | 18533 | 15.98 |
| | Anterior insula | Right | 28 | -48 | 10 | 453 | 8.19 |
| | Calcarine sulcus | Right | 28 | -48 | 10 | 1274 | 7.83 |
| | Fusiform and inferior temporal gyri | Left | -44 | -56 | -14 | 373 | 7.06 |
| | Anterior cerebellar lobe | Bilateral | 9 | -58 | -4 | 422 | 5.52 |
| | Calcarine sulcus | Left | -18 | -56 | 2 | 277 | 5.00 |
| **Target** | MDN, ventral visual stream, motor and somatosensory networks, thalamus, basal ganglia (putamen, caudate, globus pallidus) | Bilateral | 38 | -50 | -16 | 61189 | 28.31 |

*Note:* The table shows, for each contrast (i.e., paradigm epoch) and significant cluster, the labels of the regions involved, the MNI coordinates (X, Y, Z), the cluster's voxel extent and the peak *t* value. MDN: Multiple Demand Network.

**Supplementary Table 2.**

Results from decoding whole-brain searchlight analysis for each task component

| Contrast | Regions | Hemisphere | X | Y | Z | k | *t* value | $p_{FWE}$ |
|---|---|---|---|---|---|---|---|---|
| **Task Demand** | MDN, medial prefrontal cortex, inferior, middle and superior temporal gyri, precuneus, ventral visual stream, motor and somatosensory networks, basal ganglia (putamen, caudate, globus pallidus) | Bilateral | -20 | -66 | 50 | 100879 | 11.53 | <.001 |
| **Target Category** | *No clusters survived the FWE correction* | | | | | | | |
| **Relevant Feature** | Middle and inferior temporal gyri | Left | -46 | -56 | -2 | 567 | 5.44 | .025 |
| | V1 and V2 | Left | -20 | -80 | -10 | 1142 | 4.75 | .002 |

*Note:* The table shows for each contrast (i.e., task component) and significant cluster, the labels of the regions involved, the MNI coordinates (X, Y, Z) in mm, the cluster's voxel extent (k), the peak *t* value, and the FWE-corrected peak. MDN: Multiple Demand Network.

**Supplementary Table 3**.

Results from the Cross-Condition Generalization Performance (CCGP) analyses embedded into a leave-one-run-out cross-validation procedure.

| | ***Task demands*** | | | | ***Target category*** | | | | ***Relevant feature*** | | | |
|---|---|---|---|---|---|---|---|---|---|---|---|---|
| | CCGP | *z* | *p* | $p_{corr}$ | CCGP | *z* | *p* | $p_{corr}$ | CCGP | *z* | *p* | $p_{corr}$ |
| preSMA/ACC | 11.12 | 5.21 | <.001 | **<.001** | -1.90 | -1.80 | .964 | ~1 | -2.01 | -1.66 | .957 | ~1 |
| L_aI/fO | 6.98 | 4.70 | <.001 | **<.001** | -1.71 | -0.96 | .832 | ~1 | -1.66 | -1.47 | .929 | ~1 |
| R_aI/fO | 4.17 | 3.04 | .001 | **.001** | -1.32 | -1.33 | .908 | ~1 | 0.22 | 0.36 | .361 | ~1 |
| L_IFS | 6.18 | 3.85 | <.001 | **<.001** | -0.92 | -0.91 | .820 | ~1 | 1.77 | 1.43 | .076 | .686 |
| R_IFS | 7.02 | 4.59 | <.001 | **<.001** | -1.70 | -1.42 | .922 | ~1 | -0.64 | -0.58 | .719 | ~1 |
| L_IFJ | 19.54 | 5.00 | <.001 | **<.001** | -3.15 | -2.59 | .995 | ~1 | 0.46 | 0.32 | .375 | ~1 |
| R_IFJ | 8.25 | 4.61 | <.001 | **<.001** | 0.04 | 0.04 | .483 | ~1 | -0.87 | -0.86 | .483 | ~1 |
| L_IPS | 13.93 | 5.44 | <.001 | **<.001** | -2.45 | -2.53 | .994 | ~1 | -0.07 | 0.03 | .486 | ~1 |
| R_IPS | 14.34 | 5.44 | <.001 | **<.001** | -0.24 | 0.00 | .500 | ~1 | -1.27 | -1.18 | .882 | ~1 |

*Note:* The table shows the mean CCGP score for each task component and ROI after averaging across the eight cross-validation iterations, the outputs (Z statistic, *p*-value) of the one-sided, one-sample Wilcoxon signed rank test addressing above-chance CCGP scores, and the adjusted *p*-values after a Bonferroni-Holm multiple comparison correction.

**Supplementary Table 4.**

Results from CCGP whole-brain searchlight analysis for each task component

| Contrast | Regions | Hemisphere | X | Y | Z | k | *t* value | $p_{FWE}$ |
|---|---|---|---|---|---|---|---|---|
| **Task Demand** | MDN, orbitofrontal cortex, inferior, middle and superior temporal gyri, precuneus, ventral visual stream, motor and somatosensory networks, putamen. | Bilateral | -46 | -4 | 46 | 92092 | 10.20 | <.001 |
| **Target Category** | *No clusters survived the FWE correction* | | | | | | | |
| **Relevant Feature** | *No clusters survived the FWE correction* | | | | | | | |

*Note:* The table shows for each contrast (i.e., task component) and significant cluster, the labels of the regions involved, the MNI coordinates (X, Y, Z) in mm, the cluster's voxel extent (k), the peak *t* value, and the FWE-corrected peak. MDN: Multiple Demand Network.

**Supplementary Table 5.**

Results from the model-based Representational Similarity Analysis (RSA) separated by task demand, based on cross-validated Pearson correlation distances (from run-wise LSU beta maps).

| | Task demand condition | Target category | | | | Relevant feature | | | | Conjunction | | | |
|---|---|---|---|---|---|---|---|---|---|---|---|---|---|
| | | beta | *z* | *p* | $p_{corr}$ | beta | *z* | *p* | $p_{corr}$ | beta | *z* | *p* | $p_{corr}$ |
| preSMA/ACC | SEL | -0.0001 | 0.80 | 0.211 | ~1 | 0.0003 | 0.54 | 0.296 | 0.775 | -0.0011 | 0.10 | 0.458 | ~1 |
| | INT | -0.0029 | -1.29 | 0.902 | ~1 | -0.0019 | -1.30 | 0.904 | ~1 | -0.0012 | 0.43 | 0.335 | ~1 |
| L_aI/fO | SEL | -0.0020 | -0.38 | 0.649 | ~1 | 0.0013 | 0.65 | 0.258 | 0.962 | -0.0007 | -0.37 | 0.644 | ~1 |
| | INT | 0.0002 | -0.91 | 0.820 | ~1 | -0.0017 | -0.52 | 0.700 | ~1 | 0.0034 | 0.97 | 0.166 | ~1 |
| R_aI/fO | SEL | -0.0036 | -0.77 | 0.781 | ~1 | 0.0000 | 0.05 | 0.481 | 0.591 | 0.0035 | 0.24 | 0.404 | ~1 |
| | INT | -0.0025 | -0.44 | 0.670 | ~1 | 0.0023 | -0.29 | 0.613 | ~1 | 0.0011 | 0.41 | 0.340 | ~1 |
| L_IFS | SEL | 0.0045 | 0.83 | 0.203 | ~1 | 0.0062 | 1.21 | 0.114 | 0.796 | -0.0069 | -0.97 | 0.834 | ~1 |
| | INT | 0.0034 | 1.86 | 0.031 | 0.281 | 0.0051 | 1.90 | 0.028 | 0.256 | -0.0063 | -1.15 | 0.875 | ~1 |
| R_IFS | SEL | 0.0004 | 0.19 | 0.425 | ~1 | 0.0035 | 1.30 | 0.096 | 0.768 | -0.0076 | -0.50 | 0.690 | ~1 |
| | INT | 0.0023 | -0.34 | 0.634 | ~1 | -0.0066 | -1.95 | 0.974 | ~1 | 0.0037 | 1.04 | 0.149 | ~1 |
| L_IFJ | SEL | 0.0010 | 0.77 | 0.219 | ~1 | 0.0016 | 1.40 | 0.080 | 0.723 | -0.0053 | -0.94 | 0.827 | ~1 |
| | INT | 0.0031 | 0.82 | 0.207 | ~1 | 0.0019 | 1.18 | 0.119 | 0.953 | -0.0034 | -0.41 | 0.660 | ~1 |

| | | | | | | | | | | | | | |
|---|---|---|---|---|---|---|---|---|---|---|---|---|---|
| R_IFJ | SEL | 0.0052 | 1.51 | 0.065 | 0.52 | 0.0028 | 0.70 | 0.240 | 0.962 | -0.0055 | -1.56 | 0.940 | ~1 |
| | INT | -0.0040 | -1.01 | 0.844 | ~1 | -0.0069 | -1.96 | 0.975 | ~1 | 0.0112 | 1.70 | 0.045 | 0.405 |
| L_IPS | SEL | 0.0017 | 0.54 | 0.296 | ~1 | 0.0007 | 0.97 | 0.166 | 0.896 | 0.0014 | 0.09 | 0.464 | ~1 |
| | INT | -0.0004 | -0.52 | 0.700 | ~1 | 0.0000 | -0.57 | 0.714 | ~1 | -0.0024 | -0.12 | 0.547 | ~1 |
| R_IPS | SEL | 0.0057 | 1.81 | 0.035 | 0.318 | 0.0019 | 1.04 | 0.149 | 0.896 | -0.0043 | -0.91 | 0.820 | ~1 |
| | INT | 0.0009 | -0.03 | 0.514 | ~1 | -0.0039 | -1.00 | 0.841 | ~1 | 0.0011 | 1.00 | 0.159 | ~1 |

*Note:* The table shows the mean beta weights for each RSA model and ROI per task demand condition (selection and integration), the outputs (Z statistic, *p*-value) of the one-sided, one-sample Wilcoxon signed rank test addressing above-zero regression coefficients, and the adjusted *p*-values after a Bonferroni-Holm multiple comparisons correction.

**Supplementary Table 6.**

Association of task demand decoding accuracy with selection-integration behavioral differences

| | ***Behavioral Accuracy*** | | | | ***Reaction Times*** | | | |
|---|---|---|---|---|---|---|---|---|
| | r | $p$ | $p_{corr}$ | $BF_{01}$ | r | $p$ | $p_{corr}$ | $BF_{01}$ |
| preSMA/ACC | -0.155 | 0.346 | ~1 | *5.162* | -0.002 | 0.989 | ~1 | *8.018* |
| L_aI/fO | -0.332 | 0.039 | 0.35 | 0.964 | 0.089 | 0.589 | ~1 | *6.936* |
| R_aI/fO | -0.151 | 0.359 | ~1 | *5.276* | -0.007 | 0.965 | ~1 | *8.011* |
| L_IFS | 0.088 | 0.594 | ~1 | *6.962* | -0.063 | 0.705 | ~1 | *7.467* |
| R_IFS | 0.223 | 0.173 | ~1 | *3.192* | -0.233 | 0.154 | ~1 | 2.919 |
| L_IFJ | -0.107 | 0.516 | ~1 | *6.5* | 0.002 | 0.991 | ~1 | *8.018* |
| R_IFJ | 0.142 | 0.39 | ~1 | *5.555* | 0.102 | 0.536 | ~1 | *6.631* |
| L_IPS | -0.321 | 0.046 | 0.368 | 1.112 | 0.307 | 0.057 | 0.513 | 1.328 |
| R_IPS | -0.223 | 0.172 | ~1 | *3.172* | 0.085 | 0.605 | ~1 | *7.02* |

*Note:* Pearson's correlation was used to assess the relationship between subject-wise mean Task Demand decoding accuracy and subject-wise mean behavioral differences between Integration and Selection, computed separately for accuracy and reaction times. Holm–Bonferroni correction was applied within each behavioral measure to account for multiple comparisons across ROIs. For each component and ROI, the table also includes Bayes factors supporting the null hypothesis ($BF_{01}$) from Bayesian correlation analyses, quantifying evidence for the absence of an association between decoding performance and the behavioral measure (moderate to strong evidence for the null effect in *cursive*).
